# Characterization of radiations-induced genomic structural variations in *Arabidopsis thaliana*

**DOI:** 10.1101/2024.07.25.605065

**Authors:** Salimata Ousmane Sall, Abdelmalek Alioua, Sébastien Staerck, Stéfanie Graindorge, Michel Pellicioli, Jacky Schuler, Quentin Raffy, Marc Rousseau, Jean Molinier

## Abstract

DNA, is assaulted by endogenous and exogenous agents that lead to the formation of damage. In order to maintain genome integrity DNA repair pathways must be efficiently activated to prevent mutations and deleterious chromosomal rearrangements. Conversely, genome flexibility is also necessary to allow genetic diversity and evolution. The antagonist interaction between maintenance of genome integrity and flexibility determines genome shape and organization. Therefore, it is of great interest to understand how the whole linear genome structure behaves upon formation and repair of DNA damage. For this, we used long reads sequencing technology to identify and to characterize genomic structural variations (SV) of wild-type *Arabidopsis thaliana* somatic cells exposed either to UV-B, to UV-C or to protons irradiations. We found that genomic regions located in heterochromatin a more prone to form SVs than those located in euchromatin, highlighting that genome stability and flexibility differs along the chromosome. This holds true in Arabidopsis plants deficient for the expression of master regulators of the DNA Damage Response (DDR), ATM and ATR, suggesting that independent and alternative surveillance processes exist to maintain integrity in genic regions. Finally, the analysis of the radiations-induced deleted regions allowed determining that exposure UV-B, UV-C and protons induced the Microhomology-mediated end joining mechanism (MMEJ) and that both ATM and ATR repress this repair pathway.

## INTRODUCTION

DNA is the support of the genetic information and defines the genome of living organisms. Genome is organized through the linear order of genetic entities such as protein coding genes (PCG), intergenic regions (IR), transposable elements (TE) and repeats. Genome size results from the antagonist interactions between processes regulating its stability and its flexibility (Schubert and Vu, 2016). Indeed, genome stability is necessary to ensure integrity whilst genome flexibility allows genetic diversity and evolution (Melamed-Bessudo et al., 2016; Bhadra et al., 2023). Variations in genome size and organization occur through changes in ploidy level (*i.e.*, whole genome duplication) or during stress- or development-induced structural variations (SV; Schubert and Vu, 2016; Masuda et al., 2017; Zhang et al., 2023). All these changes contribute to the reshaping of the genome which occurs with different dynamics. Indeed, particular genomic regions display higher structural variability, such as centromeres containing large amounts of repeats and TE (Lian et al., 2024; Naish and Henderson, 2024).

Plants, like most of the living organisms, are exposed to numerous environmental cues that can generate point mutations and alter genome structure. The release of TE silencing can lead to their mobilization and to transposition events affecting genome organization and structure (Roquis et al., 2021; Zhang et al., 2023). Endogenous agents (*i.e.*, secondary metabolites) and exposure to environmental cues (*i.e.*, high light) induce the formation of DNA damage (Mehta and Haber, 2014; Molinier, 2017; Yousefzadeh et al., 2021). In order to maintain genome integrity and to prevent deleterious chromosomal rearrangements several different processes are activated. Transcriptional and post-transcriptional gene silencing, restrict TE mobilization, and DNA repair pathways allow maintenance of genome integrity from single nucleotide to dozens of base pairs (Bergis-Ser et al., 2024; Herbst et al., 2024). The DNA Damage Response (DDR) is orchestrated by 2 phosphatidyl-inositol 3-kinase-like (PI3) protein kinases: ATR (Ataxia-telangiectasia-mutated and Rad3-related) and ATM (Ataxia-telangiectasia-mutated; Nisa et al., 2019). ATM is the main DNA Double Strand Break (DSB) signal transducer (Szurman-Zubrzycka et al., 2023; Bergis-Ser et al., 2024; Herbst et al., 2024) whereas ATR triggers DDR during replication stress, exposure to UV and to DNA modifying agents (Szurman-Zubrzycka et al., 2023). ATM and ATR phosphorylate specific factors albeit they also share some common targets (Shiloh, 2001; Roitinger et al., 2015). ATM and ATR deficient Arabidopsis plants exhibit hypersensitivity to DSB-and replicative stress-inducing agents, respectively (Culligan et al., 2006; Garcia et al., 2000, 2003). Although ATM mutant plants are sterile (Garcia et al., 2003), the combination of both mutations allows the recovery of plants (Vespa et al., 2005).

DSBs are induced upon exposure to genotoxic agents (*i.e.*, non-ionizing and ionizing radiations) and to biotic/abiotic stresses (Lucht et al., 2002; Kovalchuk et al., 2003; Molinier et al., 2005; Mehta and Haber, 2014). DSBs are also formed by repair intermediates of different types of damage (Sobol et al., 2003; Reitz, et al., 2023), during replication (Schuermann et al., 2009; Waterworth et al., 2011), during transpositions events (Hedges and Deininger, 2007) and during meiosis by the specific endonuclease SPO11 (Grelon et al., 2001). DSB is the most deleterious DNA damage and thus must be efficiently repaired. DSBs are processed by 2 main pathways: Non-Homologous End-Joining (NHEJ) and Homologous Recombination (HR; Puchta, 2005; Schuermann et al., 2005). NHEJ is an error prone process that ligates break ends with most of the time a loss of genetic information (Puchta, 2005). Conversely, HR is an accurate repair mechanism that uses a homologous sequence found on the sister chromatid or within the homologous chromosome (Puchta, 2005). The predominant DSB repair pathway used in plants is NEHJ (Puchta, 2005). Other DSB repair pathways rely on homology-directed repair or DNA synthesis (Puchta, 2005; Schubert and Vu, 2016). The Microhomology-mediated end joining (MMEJ), involves alignment of micro-homologous sequences (2 to 20 bp) in the vicinity of the DNA break and leads to variable sizes of deletions (Puchta, 2005; Schubert and Vu, 2016). The Synthesis-dependent strand-annealing (SDSA) pathway uses homologous DNA templates by strand displacement (Puchta, 2005; Schubert and Vu, 2016). SDSA allows mostly error free repair although insertions originating from various sequences templates could occur (Puchta, 2005).

In somatic plant cells, several different approaches have been developed to monitor the use of these different DSB repair pathways. Exogenous templates (*i.e.*, linearized plasmids; Puchta and Hohn, 1991; Dubest et al., 2002; Orel and Puchta, 2003), transgenes (Swodoba et al., 1994; Molinier et al., 2004), CRISPR Cas9 combined with high throughput short reads sequencing revealed how NHEJ/HR are used and led to the identification of various chromosomal rearrangements (Vu et al., 2017; Samach et al., 2023). Indeed, these strategies allowed the characterization of the DSB repair pathway choice (Vu et al., 2017 and of the factors involved, directly or indirectly, in the different repair processes (Vu et al., 2017). We highly gained in knowledge with the use of CRISPR Cas9 to induce DNA breaks within particular genomic/epigenomic contexts to characterize the outcome of repair (Vu et al., 2017). Nevertheless, it remains to be deciphered how the whole linear genome organization behaves upon induction of different types of DNA damage. Indeed, DNA damage formation and the choice of the DNA repair may vary according to the genomic and epigenomic contexts (*i.e*, chromatin compaction level; Johann to Berens and Molinier, 2020) and thus could influence the balance between genome stability and flexibility (Johann to Berens and Molinier, 2020). The genome wide analysis of structural variations (SV) would provide an overview of the chromosomal rearrangements that may have occurred in each genomic/epigenomic contexts independently of a targeted sequence. This is made possible with the development of the third-generation sequencing methods producing long reads (> 50 kb; Marx, 2023) and allowing an accurate coverage of the genome, including repetitive sequences (Naish et al., 2021, 2024). This paves the way for the genome wide identification and characterization of SV.

In this study we addressed the question of the induction of genomic SV in somatic Arabidopsis plant cells upon exposure to ionizing (protons) and non-ionizing radiations (UV-B and UV-C). Using long reads sequencing technology, we first characterized, in 3 independent biological replicates, the putative SV of WT Arabidopsis plants originating from our collection of seeds in comparison with the publicly available reference genome TAIR10 (https://www.arabidopsis.org/). This allowed defining the pedigree of our plants to further characterize, quantitatively and quantitatively, the repertoire of SV, induced in WT Arabidopsis plants exposed to UV-B, UV-C and protons. We identified that ionizing and non-ionizing radiations triggered to same types of chromosomal rearrangements and that constitutive heterochromatin is more prone to form these SV than other part of the (epi)genome.

We also analyzed the linear genome structure of Arabidopsis plants deficient for the expression of ATM and ATR, the master regulators of the DDR. We found that euchromatic genic regions are less prone to be rearranged than heterochromatic TE-IR, highlighting that different and complex genome surveillance mechanisms exist along the genome, even in the absence of the main DDR factors. Finally, the long reads sequencing data allowed determining to which extents, NHEJ and homology-directed DSB repair have been used. We characterized radiation-induced MMEJ events and identified that ATM and ATR repress this repair pathway.

This study provides the genome wide profiles of rearrangements induced upon exposure to non-ionizing and ionizing radiations and documents that genome stability and flexibility differ along the chromosome territories.

## RESULTS

### Experimental design

In order to characterize genotoxic stress-induced SV in different Arabidopsis genotypes we needed to define to which genomic data we had to refer to. First, we characterized whether WT Arabidopsis plants originating from our collection of seeds and used as untreated control for each type of treatment, contained genomic SV such as insertions, deletions, duplications, inversions and inversions duplications, relative to the TAIR10 reference genome (https://www.arabidopsis.org/; Fig. 1A). For this, we performed long reads sequencing of the genomic DNA prepared from 3 independent biological replicates of untreated WT (Col-0) Arabidopsis plants. Importantly, these biological replicates correspond to the untreated control of each genotoxic treatment (Fig. 1A). This allowed defining our plant pedigree (Fig. 1A). Second, we identified the radiations-induced SVs by sequences comparison with the TAIR10 reference genome and upon subtraction of the total SVs found in our pedigree (Fig. 1B). This allowed the identification of SV for each source of radiations (Fig. 1B). Third, we determined SV in DDR mutant plants using long reads sequencing data from *atm*, *atr* and *atm atr* untreated plants compared with the TAIR10 reference genome and with our pedigree. This allowed determining *atm*, *atr* and *atm atr* SVs (Fig. 1C). Lastly, we identified the radiations-induced SV in both *atm*- and *atr*-treated plants relative to the TAIR10 reference genome and upon subtraction with the SV found in the corresponding untreated mutant and in WT treated plants (Fig. 1D and 1E). This experimental design is thought to take into account SV originating from all genetic backgrounds (WT and DDR deficient plants) in order to identify SV induced by the different sources of radiations.

**Figure 1:**
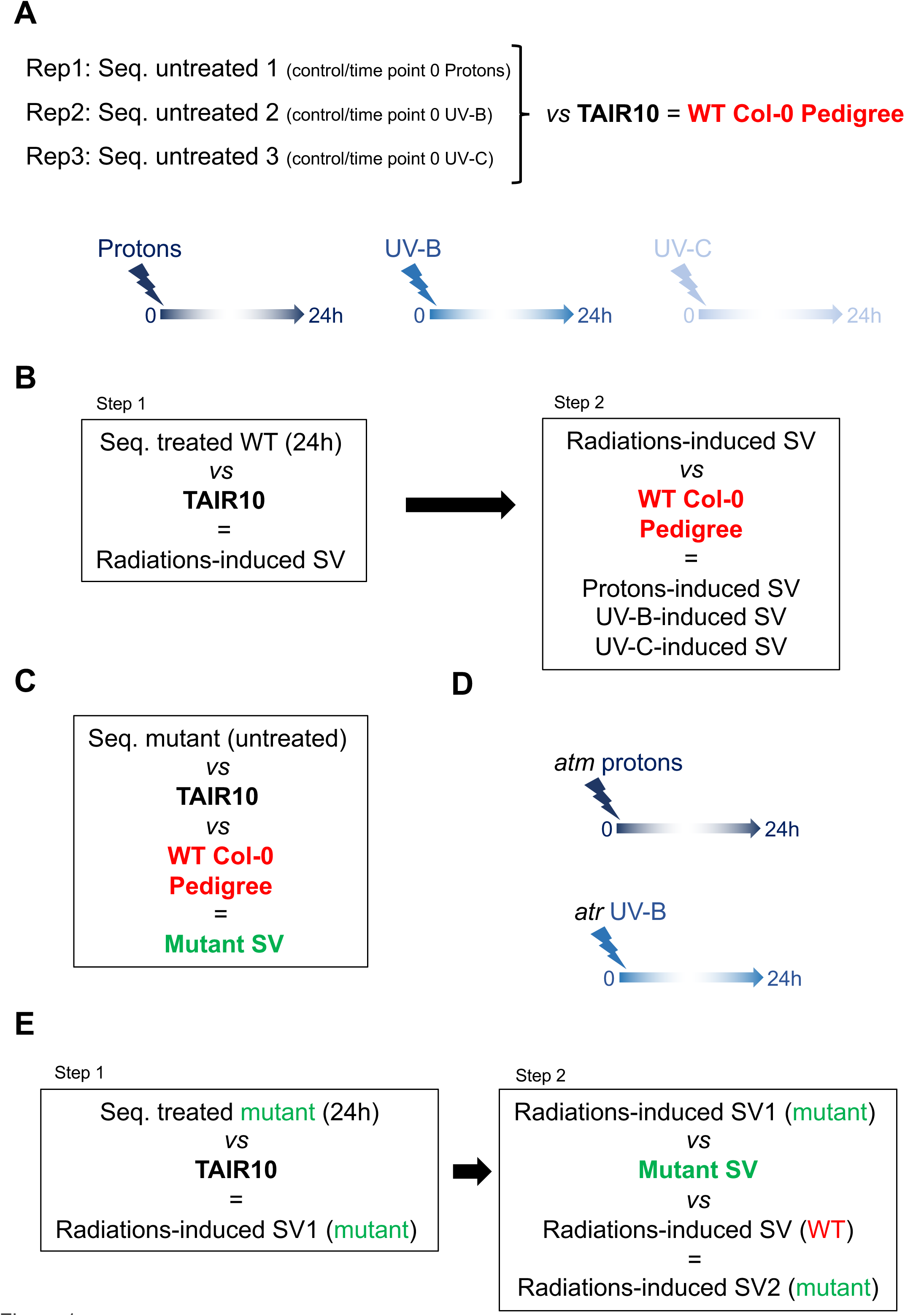
Experimental design. **A.** WT, Col-0, Arabidopsis plants were irradiated with either protons, UV-B or UV-C. 24h upon each treatment, plant material was harvested to determine radiations-induced SV using long reads sequencing. Each untreated control (time point 0) of the 3 different types of irradiations, have been used to characterize the pedigree of our collection of WT (Col-0) Arabidopsis seeds. The comparison of the genomic sequences (seq.) with the TAIR10 reference genome allows defining the SVs of the plants originating from our collection of WT (Col-0) Arabidopsis seeds. The total number of SV defines our pedigree and is thus used as reference for further experiments. **B.** Schematic representation of the approach designed to characterize UV-B-, UV-C- and protons-induced SV in WT plants. Step 1: long reads sequences (seq.) of treated plants are compared to the TAIR10 reference genome to identify radiations-induced SV. Step 2: SVs of our pedigree are subtracted to radiations-induced SV to determine the UV-B-, UV-C- and protons-induced SVs. **C.** Schematic representation of the approach designed to characterize SV in DDR deficient plants. Long reads sequences (seq.) of untreated *atm* and *atr* plants are compared to the TAIR10 reference genome to identify SV in mutant plants. **D.** *atm* and *atr*, plants were irradiated with protons and UV-B, respectively. 24h upon each treatment, plant material was harvested to determine radiations-induced SV using long reads sequencing. **E.** Same as **B.** for *atr* UV-B- and *atm* protons-treated plants.

### Characterization of WT plants pedigree

In order to characterize the SVs that may contain the somatic cells of WT (Col-0) untreated plants, we compared the long reads sequencing data obtained from 3 independent biological replicates with the TAIR10 reference genome. In each replicate we identified 229, 339 and 291 SV, such as insertions, deletions, duplications, inversions and inversions-duplications (Fig. 2A). The distribution of the different types of SV does not vary between replicates (Fig. 2A). INDELS (insertions-deletions) are predominantly represented (> 80%; Fig. 2A) and their sizes do not significantly differ between replicates (Fig. 2B) suggesting a limited variability among samples.

**Figure 2:**
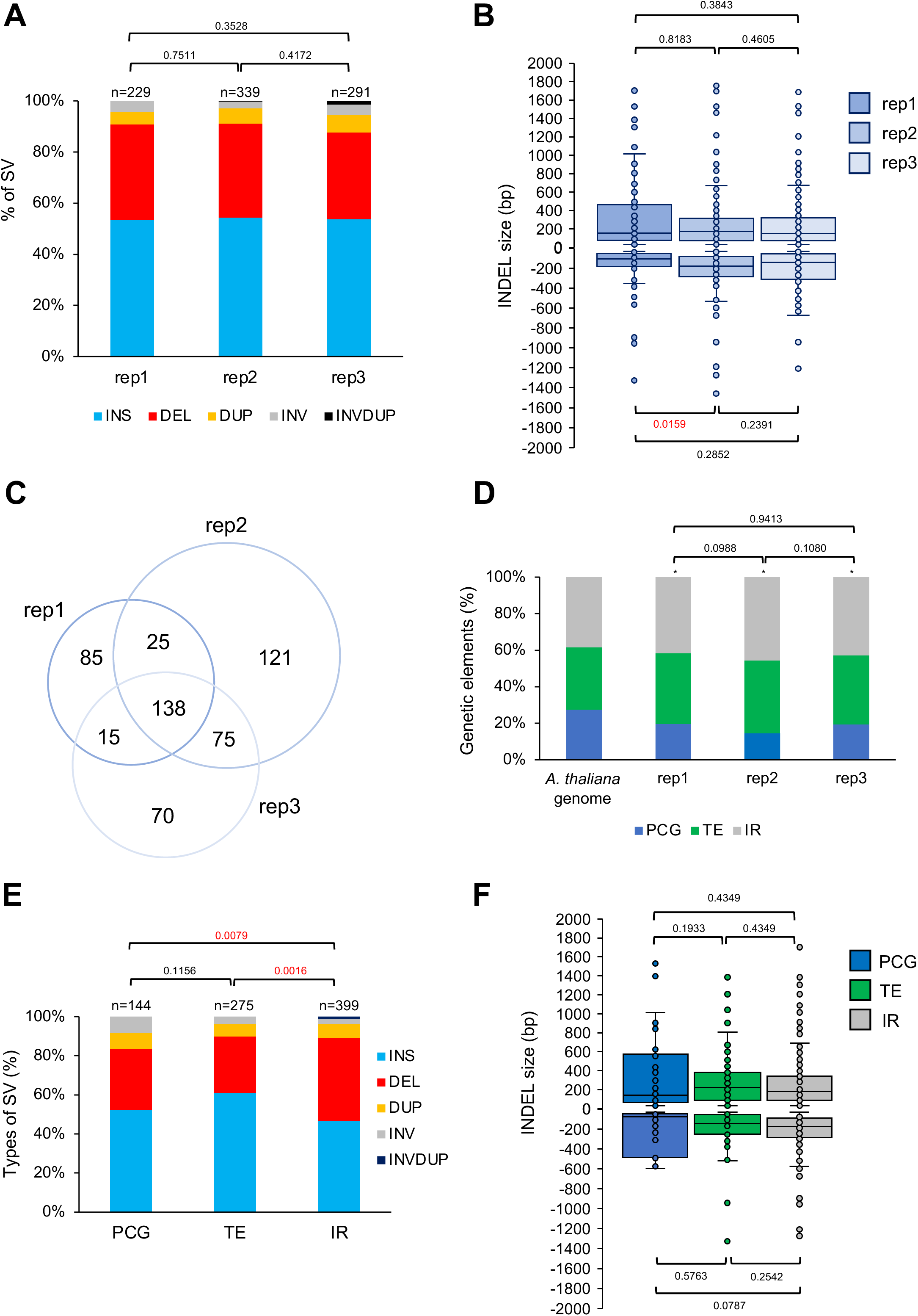
Characterization of the structural variations in WT Arabidopsis plants. **A.** Histogram representing the distribution of the different types of genomic SV identified in each of the 3 independent biological replicates of WT (Col-0) Arabidopsis plants. INS: Insertion; DEL: Deletion; DUP: Duplication; INV: Inversion; INVDUP: Inversion Duplication. n= total number of SV. Exact p values are shown (Chi square test). **B.** Box plots representing the size of the INDELs identified in each of the 3 independent biological replicates. Exact p values are shown (Mann Whitney Wilcoxon test). **C.** Venn diagram representing the overlap of SV between the 3 independent biological replicates. **D.** Histogram representing the distribution of the genetic elements (Protein Coding Genes: PCG; Transposable Elements: TE and Intergenic regions: IR) exhibiting SV in each of the 3 independent biological replicates. The distribution of the genetic elements in the *A. thaliana* genome is shown. Exact p values are shown, * p<0.01 *vs A. thaliana* genome (Chi square test). **E.** Histogram showing the distribution of the genomic SV (INS: Insertion; DEL: Deletion; DUP: Duplication; INV: Inversion; INVDUP: Inversion Duplication) identified in the genetic elements (Protein Coding Genes: PCG; Transposable Elements: TE and Intergenic regions: IR). Exact p values are shown (Chi square test). **F.** Box plots representing the INDELs sizes identified in the genetic elements (Protein Coding Genes: PCG; Transposable Elements: TE and Intergenic regions: IR). Exact p values are shown (Mann Whitney Wilcoxon test).

The comparison of the SV found in each replicate shows that 138 are common between the 3 samples, 7% to 22% are shared between 2 replicates and 27 to 37% are unique to each replicate (Fig. 2C). This shows that a core of SV exists in the Arabidopsis plants originating from our collection of seeds but also that some SV are present at lower frequencies.

The distribution of the genetic elements: Protein Coding Genes (PCG); Transposable Elements (TE) and Intergenic regions (IR), affected by SV are not significantly different between replicates (Fig. 2D). Importantly, TE and IR represent more than 80% of the location of the SV (Fig. 2D) stressing that these genetic entities are more prone to be rearranged than PCG. Deletions occur more often in IR than in TE and PCG but their sizes remain similar between genetic entities (Fig. 2E and 2F). The overrepresentation of TE and IR exhibiting SV, suggests that, particular genetic elements might preferentially undergo rearrangements and that particular genomic/epigenomic features might exist.

Thus, we compared the location of the identified SV with the coordinates of centromeres/chromosomes arms and with the publicly available Arabidopsis epigenomic landscape defined as chromatin states (CS; Sequeira-Mendes et al., 2014). Most of the SV (>70%) overlap with centromeric and pericentromeric regions (Fig. 3A) and with the repressive chromatin states: CS8 and 9 (Sequeira-Mendes et al., 2014; Fig. 3B). This highlights that constitutive heterochromatin, enriched in repetitive elements, is more prone to form SV than euchromatin. Interestingly, LTR/Gypsy and DNA TE superfamilies are significantly overrepresented among all TE containing SV (Fig. S1) indicating that these types of TE exhibit more flexibility than others.

**Figure 3:**
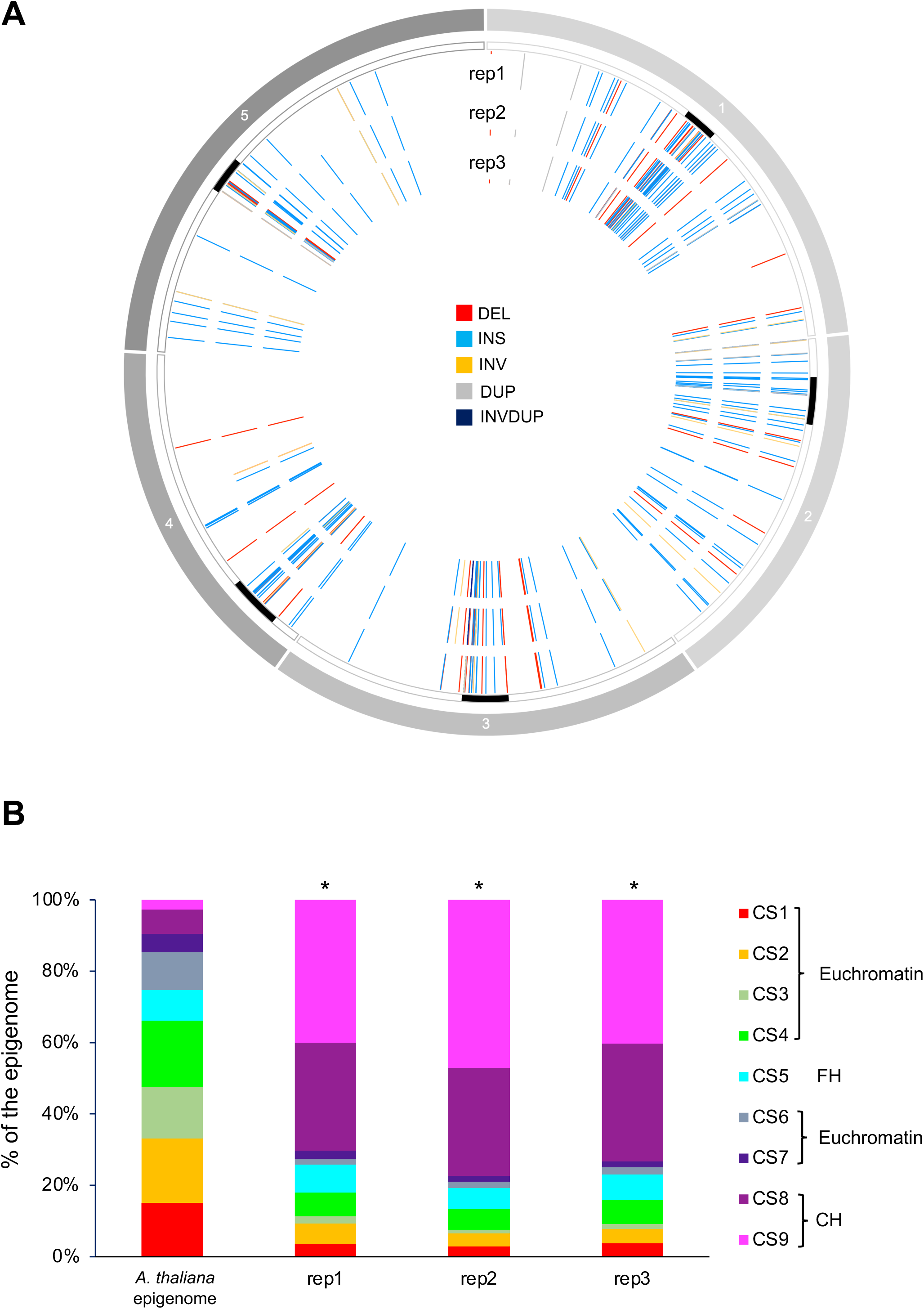
Genomic location and epigenomic features of the structural variations identified in WT Arabidopsis plants. **A.** Circos representation of genomic SV (INS: Insertion; DEL: Deletion; DUP: Duplication; INV: Inversion; INVDUP: Inversion Duplication) identified in each independent biological replicate. Black rectangles represent the centromeres. **B.** Histogram representing the distribution of the chromatin states (CS) overlapping with the SV identified in each independent biological replicates. Chi square test *< 0.01 compared to the CS distribution in the Arabidopsis epigenome (Sequeira-Mendes et al., 2014). CH: constitutive heterochromatin; FH: facultative heterochromatin.

Hence, the use of long reads sequencing allows the identification of genomic SV in Arabidopsis WT plants originating from our collection of seeds in comparison with the Arabidopsis TAIR10 reference genome. Importantly, these results indicate that genomic variability exists between plants that have been propagated and with the reference TAIR10 genome. Moreover, it shows that these changes occur predominantly in constitutive heterochromatin containing high number of repetitive sequences that are thought to be more prone to be rearranged (Lian et al., 2024; Naish and Henderson, 2024). Therefore, these genomic data will be used as references to qualitatively and quantitatively determine the effect of genotoxic stress exposure on genome integrity of WT Arabidopsis plants.

### Exposure to radiations induces structural variations predominantly in constitutive heterochromatin

The use of long reads sequencing of 3 independent biological replicates of WT Arabidopsis plants originating from our collection of seeds allowed determining their pedigree. We used this as reference, to quantitatively and qualitatively characterize genotoxic stress-induced SV. We treated WT Arabidopsis plants with non-lethal doses of either non-ionizing (UV-B, UV-C) or ionizing radiations (protons). UV-B and UV-C induce photoproducts (Molinier, 2017), DSB (Peak and Peak, 1990; Ries et al., 2000; Molinier et al., 2004) and to a lower extent oxidatively-induced DNA damage (UV-B; Cadet et al., 2015). Protons exposure mostly leads to the formation of reactive oxygen species (ROS) and to DSB (Ward, 1988; Kim et al., 2019). Importantly, the use of these sources of radiations allows an acute exposure of plants to the genotoxic agent that differs from treatments with chemicals (*i.e.*, cisplatin) which is much more related to a chronic exposure. Moreover, the kinetics of uptake of a chemical and the time window of the formation of damages, is much more difficult to define. Long reads sequencing was performed on genomic DNA prepared from samples harvested 24h upon irradiation, when DNA is thought to be repaired (Fig. 1A). Radiations-induced SVs were retrieved after subtraction of the total amount of SV found in the 3 replicates of untreated WT plants (Fig. 1B). Indeed, each replicate corresponded to the untreated control of each genotoxic treatment (Fig. 1A and 1B).

Exposure of WT Arabidopsis plants to UV-B, UV-C or protons led to the formation of 40, 64 and 49 SV, respectively, and are enriched in INDELS (Fig. 4A). Interestingly, protons treatment induced more deletions than UV-B or UV-C treatments (Fig. 4A) in relationship with the high energy delivered by ionizing radiations and their deleterious effect on DNA (Cadet et al., 2014; Ward 1988). Insertions represent more than 50% of the SV induced upon exposure to UV-B and UV-C; (Fig. 4A). This observation highlights that the outcome of repair differs upon exposure to non-ionizing and ionizing radiations likely due to the types of DNA damage induced and to the repair processes used (Molinier, 2017; Kim et al., 2019). Nevertheless, the sizes of the radiations-induced INDELs do not differ significantly among treatments (Fig. 4B). Both TE and IR are mainly affected by SV (Fig. 4C). LTR/Copia (Class I) are predominantly altered upon protons treatment whilst Class II TE (*i.e.*, MuDR) represent the main superfamily exhibiting SV upon non-ionizing radiation exposure (Fig. S2). Thus, the source of irradiation might influence the type of TE superfamily in which SV occurred, likely in relationship with their transcriptional activity and/or their intrinsic features. INDEL sizes in the different genetic entities remain invariable between treatments (Fig. S3A). Interestingly, UV-B treatment leads to a significant higher proportion of SV in PCG (mainly insertions) compared to UV-C and protons irradiations (Fig. 4C, S3B) suggesting that this genotoxic agent, preferentially induces variability in certain genetic entities.

**Figure 4:**
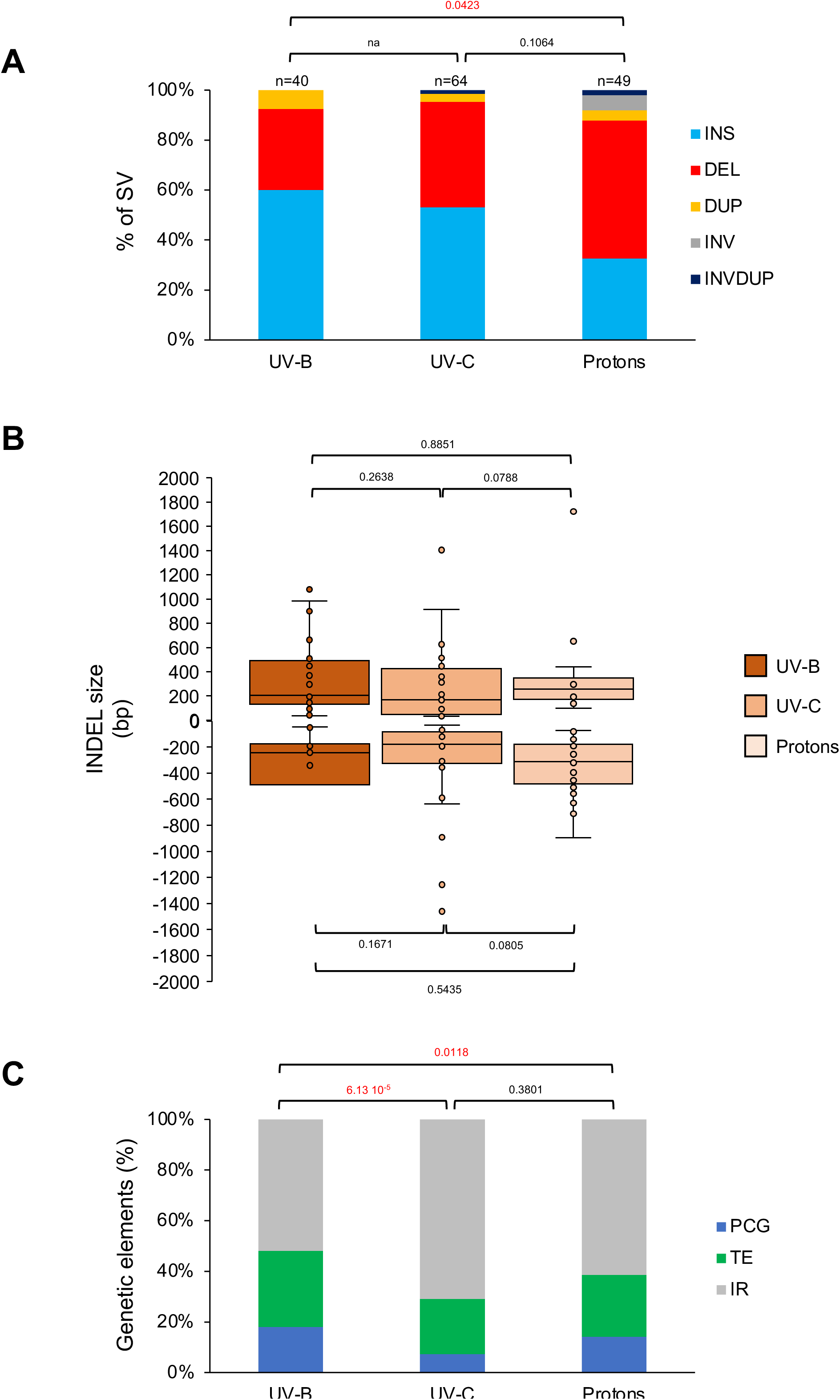
Characterization of the radiation-induced genomic structural variations in WT Arabidopsis plants. **A.** Histogram representing the distribution of the genomic SV identified in WT Arabidopsis plants treated with either UV-B, UV-C or protons. INS: Insertion; DEL: Deletion; DUP: Duplication; INV: Inversion; INVDUP: Inversion Duplication. n= total number of SV. Exact p values are shown (Chi square test). na: non-applicable **B.** Box plots representing the INDELs sizes identified in WT Arabidopsis plants treated with either UV-B, UV-C or protons. Exact p values are shown (Mann Whitney Wilcoxon test). **C.** Histogram representing the distribution of the genetic elements (Protein Coding Genes: PCG; Transposable Elements: TE and Intergenic regions: IR) exhibiting SV upon exposure to either UV-B, UV-C or protons. Exact p values are shown (Chi square test).

Given that insertions represent a good proportion of the SV (Fig. 4A), their detailed analyses would provide information about the origin of the inserted sequences. Indeed, transposition can lead to *de novo* insertions in the genome (Muñoz-López and García-Pérez, 2010). In addition, different types of sequences (within the same chromosome or between chromosomes) can be used as template for synthesis-dependent repair (*i.e.*, SDSA) and as filler DNA for repair (Gorbunova and Levy, 1997). We found that most of the inserted sequences originated from TE and IR (Fig. S4A) highlighting that these genetic entities are more prone to be used as template for insertions. However, in protons-irradiated plants we found that more insertions originated from PCG (Fig. S4A), showing that ionizing radiation would rather trigger the use of PCG regions as template for insertions. Interestingly, none of the analyzed insertions revealed a *per se* transposition event, but only truncated TE.

We also determined which genetic elements have inserted into PCG, TE and IR. Although the numbers are quite low (between 3 to 20 events), we found a trend to have insertions originating from PCG into PCG, from TE into TE and from IR into IR with a preference for an intrachromosomal origin (Fig. S4B-D and S5). These results highlight that genomic regions in the vicinity of the DNA breaks are likely used a template. This could be either due to the linear 1D organization or 3D genome structure.

We identified that constitutive heterochromatin contains much more SV than euchromatin in the untreated Arabidopsis plants (Fig. 5A). Thus, we investigated whether non-ionizing and ionizing radiations would have also triggered the formation of SV in particular chromatin contexts. In all treatments, SV are located predominantly in centromeric-pericentromeric regions overlapping with CS8 and 9 (Fig. 5B) indicating that constitutive heterochromatin is more prone to exhibit structural variability than euchromatin.

**Figure 5:**
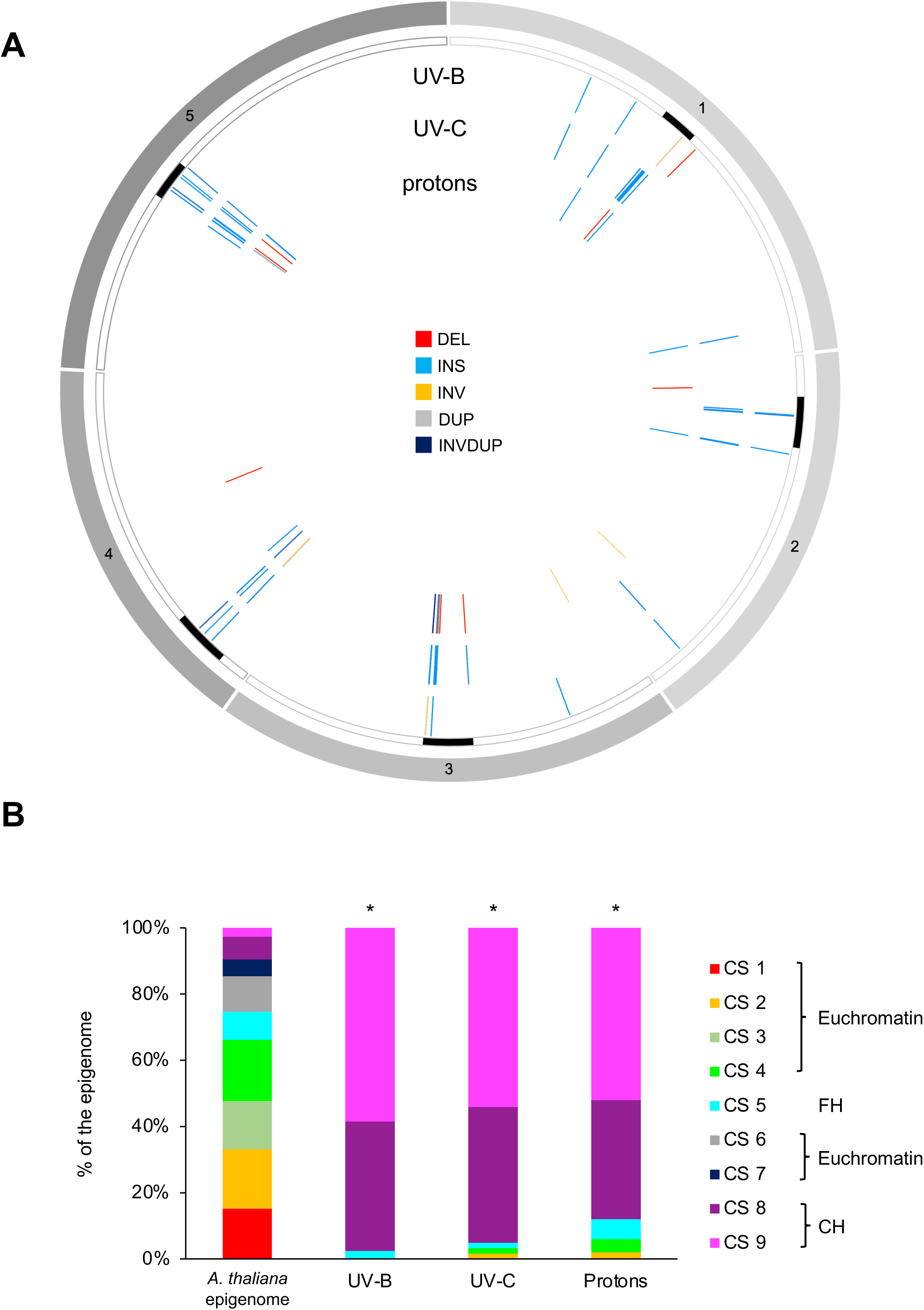
Genomic locations and epigenomic features of genomic structural variations identified in WT Arabidopsis plants irradiated with either UV-B, UV-C or protons. **A.** Circos representation of genomic SV (INS: Insertion; DEL: Deletion; DUP: Duplication; INV: Inversion; INVDUP: Inversion Duplication) identified upon exposure of WT Arabidopsis plants to either UV-B, UV-C or protons. Black rectangles represent the centromeres. **B.** Histogram representing the distribution of the chromatin states (CS) overlapping with the SV identified upon exposure to either UV-B, UV-C or protons. Chi square test *< 0.01 compared to the CS distribution in the Arabidopsis epigenome (Sequeira-Mendes et al., 2014).

The Arabidopsis genome contains regions, called hotspots of rearrangements (HOT), that would facilitate evolutionary responses to rapidly changing environmental challenges (Jiao and Schneeberger, 2020). Thus, we determined whether non-ionizing and ionizing radiations have led to the formation of SV in such genomic regions. We found 103 radiations-induced SV (> 67% of the SV), that overlap with HOT regions (Fig. S6A and B) suggesting that these genomic regions exhibiting flexibility upon exposure to environmental cues and/or genotoxic stress, may contain particular features.

Altogether, the long reads whole genome sequencing of plants exposed to different types of radiations uncovered that constitutive heterochromatin is more prone to form SV than other part of the (epi)genome and that particular regions exhibit more flexibility than others.

### Control of genome integrity by ATM and ATR

The PI3-like kinases, ATM and ATR, activates the DDR to maintain genome integrity in the face of endogenous or exogenous exposures to genotoxic agents (Shiloh, 2001). In order to better define the role of these kinases in the maintenance of genome linear structure we performed the ONT sequencing of *atm*, *atr* single mutant plants and of *atm atr* double mutant plants. In single *atr* and *atm* mutant plants, SV have been determined in comparison with the TAIR10 reference genome (Fig. 1C). In double *atm atr* mutant plants, SV have been retrieved from the comparison with the TAIR10 reference genome and with each single mutant plants (Fig. 1C).

We identified 98, 70 and 67 SV in *atm*, *atr* and *atm atr* plants, respectively (Fig. 6A). INDELS represent the most predominant types of SV (Fig. 6A). Interestingly, *atm atr* plants contain a larger proportion of INDELS compared to each single mutant plants (Fig. 6A) showing an additive effect of both mutations. Conversely, the size of INDELS do not significantly differ between single and double mutant plants (Fig. 6B) suggesting that both ATM and ATR do not regulate repair mechanisms influencing INDELS lengths but rather different repair pathways leading to INDELS. Interestingly, inversions-duplications represent more than 7% of the SV in *atm atr* plants (Fig. 6A). This type of SV represents the common process controlling copy number variation (CNV; Schubert and Vu, 2016) and suggests that error-free repair mechanisms may have been derepressed in these double mutant plants.

**Figure 6:**
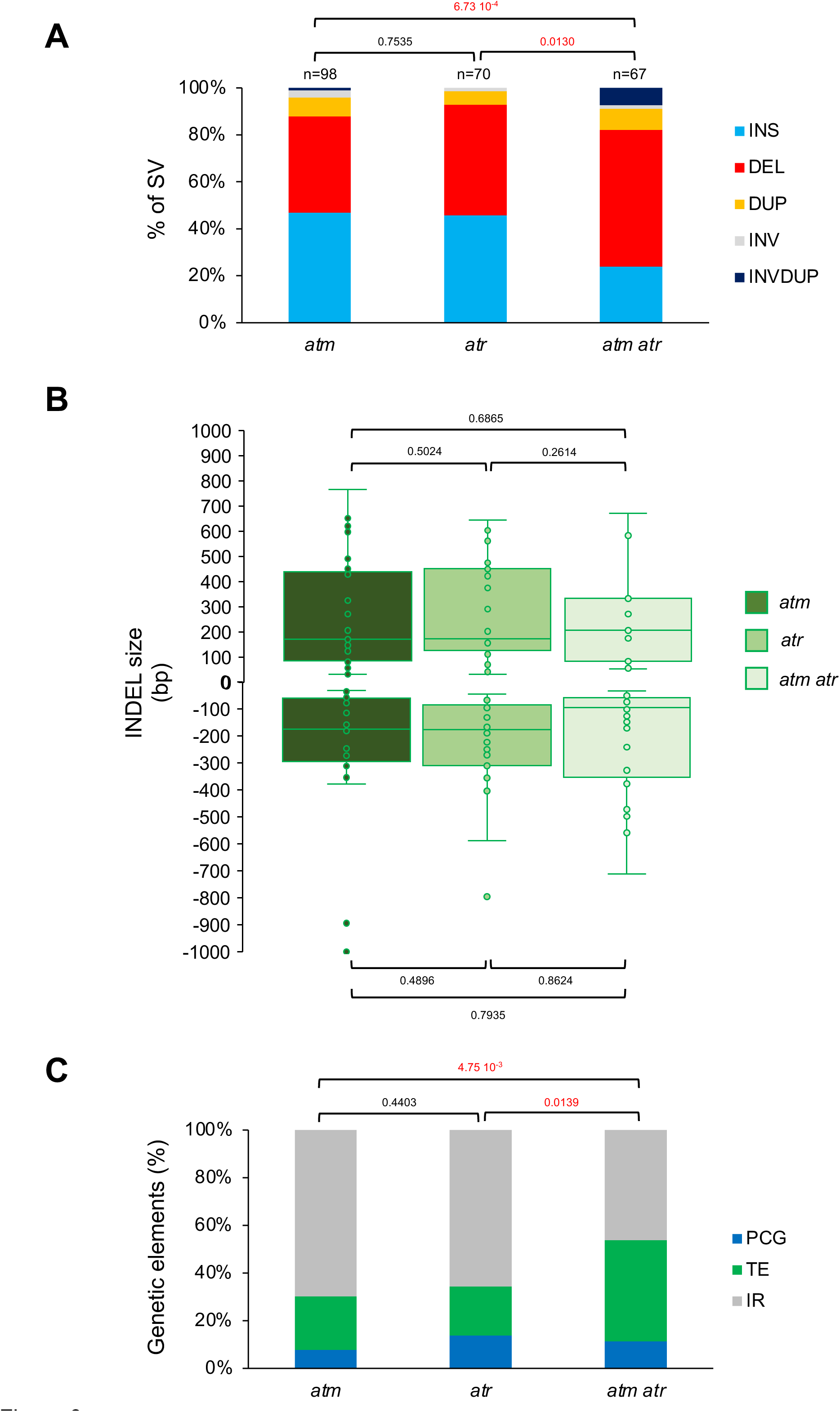
Characterization of the radiation-induced genomic structural variations in *atm*, *atr* and *atm atr* Arabidopsis plants. **A.** Histogram representing the distribution of the genomic SV identified in *atm*, *atr* and *atm atr* Arabidopsis plants. INS: Insertion; DEL: Deletion; DUP: Duplication; INV: Inversion; INVDUP: Inversion Duplication. n= total number of SV. Exact p values are shown (Chi square test). **B.** Box plots representing the INDELs sizes identified in *atm*, *atr* and *atm atr* Arabidopsis plants. No significant differences have been found between genotypes (Mann Whitney Wilcoxon test). **C.** Histogram representing the distribution of the genetic elements (Protein Coding Genes: PCG; Transposable Elements: TE and Intergenic regions: IR) exhibiting SV in *atm*, *atr* and *atm atr* Arabidopsis plants. Exact p values are shown (Chi square test).

While, in single *atm* and *atr* mutant plants, IR represent the main genomic regions exhibiting SVs, TE are more affected in *atm atr* double mutant plants (Fig. 6C). This might reflect a synergism between both PI3-like kinases in the maintenance of genome integrity in particular genetic entities. Around 75% of the SV in *atm* and *atr* plants occurred in centromeric regions highlighting that this part of the genome is, like in WT plants, more prone to be rearranged (Fig. S7A). This also suggests that likely other factors than ATM and ATR contribute to the maintenance of genome integrity in chromosomes arms.

In order to further investigate the role of ATM and ATR on genome stability, we decided to expose *atm* and *atr* plants to protons and UV-B, respectively. We did not use *atm atr* plants in this experiment due to the low recovery rate of this double mutant. Protons irradiations forms mainly DSB that are signaled by ATM (Shiloh, 2001) whilst UV-B exposure induces the formation of photodamage interfering with transcription, replication, which is preferentially signaled by ATR (Shiloh, 2001). SV previously identified in untreated *atr* and *atm* plants have been subtracted to each corresponding treated mutant plants in order to identify radiations-induced SV (Fig. 1D and 1E).

UV-B and protons irradiations induced the formation of 59 and 76 SV in *atr* and *atm* plants, respectively (Fig. 7A). Importantly, each treatment did not lead to a significant redistribution of SV types in both mutant plants (Fig. 7A) showing that irradiations did not change the outcome of the repair processes. INDELS are the predominant SV formed and their sizes remain significantly unchanged between untreated and treated mutant plants (Fig. 7B). Both TE and IR represent the majority of the genetic entities containing SV which are located in the vicinity of chromocenters (Fig. 7C and S7B). These results suggest, that even upon exposure to radiations, *ATM* or *ATR* do not regulate repair pathways leading to the formation of a particular types of SV at specific genetic entities.

**Figure 7:**
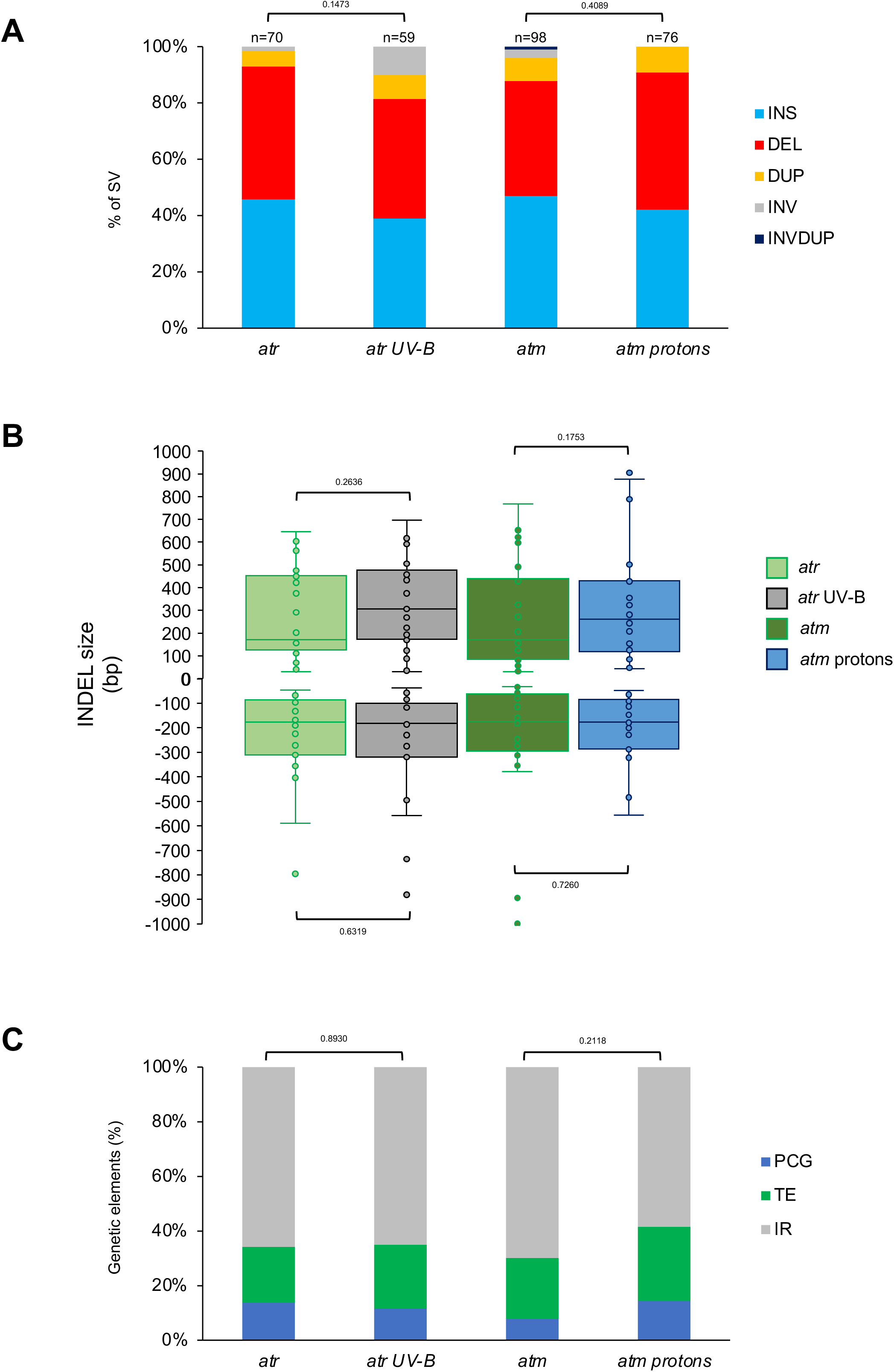
Characterization of the radiation-induced genomic structural variations in *atr* and *atm* Arabidopsis plants. **A.** Histogram representing the distribution of the genomic SV identified in *atr*, UV-B-treated *atr*, *atm* and protons-treated *atm* arabidopsis plants. INS: Insertion; DEL: Deletion; DUP: Duplication; INV: Inversion; INVDUP: Inversion Duplication. n= total number of SV. Exact p values are shown (Chi square test). **B.** Box plots representing the size of the INDELs identified in in *atr*, UV-B-treated *atr*, *atm* and protons-treated *atm* Arabidopsis plants. No significant differences have been found between untreated and treated plants (Mann Whitney Wilcoxon test). **C.** Histogram representing the distribution of the genetic elements (Protein Coding Genes: PCG; Transposable Elements: TE and Intergenic regions: IR) exhibiting SV in *atr*, UV-B-treated *atr*, *atm* and protons-treated *atm* Arabidopsis plants. Exact p values are shown (Chi square test).

We also compared SV between WT- and mutant-treated plants to uncover whether irradiation would reveal a particular role for these PI3-like kinases in the protection of genomic regions/ entities or against the formation of certain types of SV. Interestingly, we found that protons-treated *atm* plants exhibit more deletions and insertions, than WT-irradiated plants (Fig. S8A). This reveals that ATM mainly restricts INDELs formation upon protons exposure. Conversely, both UV-B-treated WT and *atr* plants display the same distribution of SVs (Fig. S8B). Irradiation and absence of the PI3-like kinases did not lead to a pronounced effect on genome integrity at particular genetic entities (Fig. S8C, D). Surprisingly, protons exposure of *atm* plants leads to significant shorter deletions than WT-treated plants (Fig. S8E, F) highlighting that particular DSB repair mechanisms might have been derepressed in this mutant plant.

Altogether, these analyses show that ATM and/or ATR regulate genome integrity mostly at TE-IR and that genic regions are less prone to be rearranged even in the absence of these PI3-like kinases. Thus, these results address the question of the putative existence of ATM-, ATR-independent DDR mechanisms in chromosomes arms (*i.e.*, at PCG).

### Analysis of genomic regions flanking the deletions highlights homology-directed DSB repair

DSB are repaired by either NHEJ or HR pathways, that are error prone and error free mechanisms, respectively (Puchta, 2005, Schuermann et al., 2005). Several mechanisms also use homologous sequences flanking the DSB to perform homology-directed DSB repair (Puchta, 2005). Interestingly, long reads sequencing data allowed determining to which extents, NHEJ and homology-directed DSB repair have been used in our different experimental conditions. For this, we analyzed the flanking regions of the full set of deletions identified in WT-treated Arabidopsis plants and in both untreated-treated *ATM* and/or *ATR* deficient plants. Importantly, we mainly identified NHEJ patterns and 1 MMEJ event in WT untreated plants among the 213 deletions analyzed (Fig. 8A). In WT-irradiated plants, between 7.5 and 23% of the genomic regions flanking directly the deletions contain homologous sequences suggesting that MMEJ repair pathway has been used (Fig. 8A and Table S1). However, the source of irradiation did not significantly change the rate of NHEJ *vs* MMEJ (Fig. 8A). The length of these micro homologies (2 and 6 bp) strongly suggests that the MMEJ pathway has been used to repair the UV-induced DNA breaks and would rather rule out the use of the SSA pathway (Fig. 8B and Table S1).

**Figure 8:**
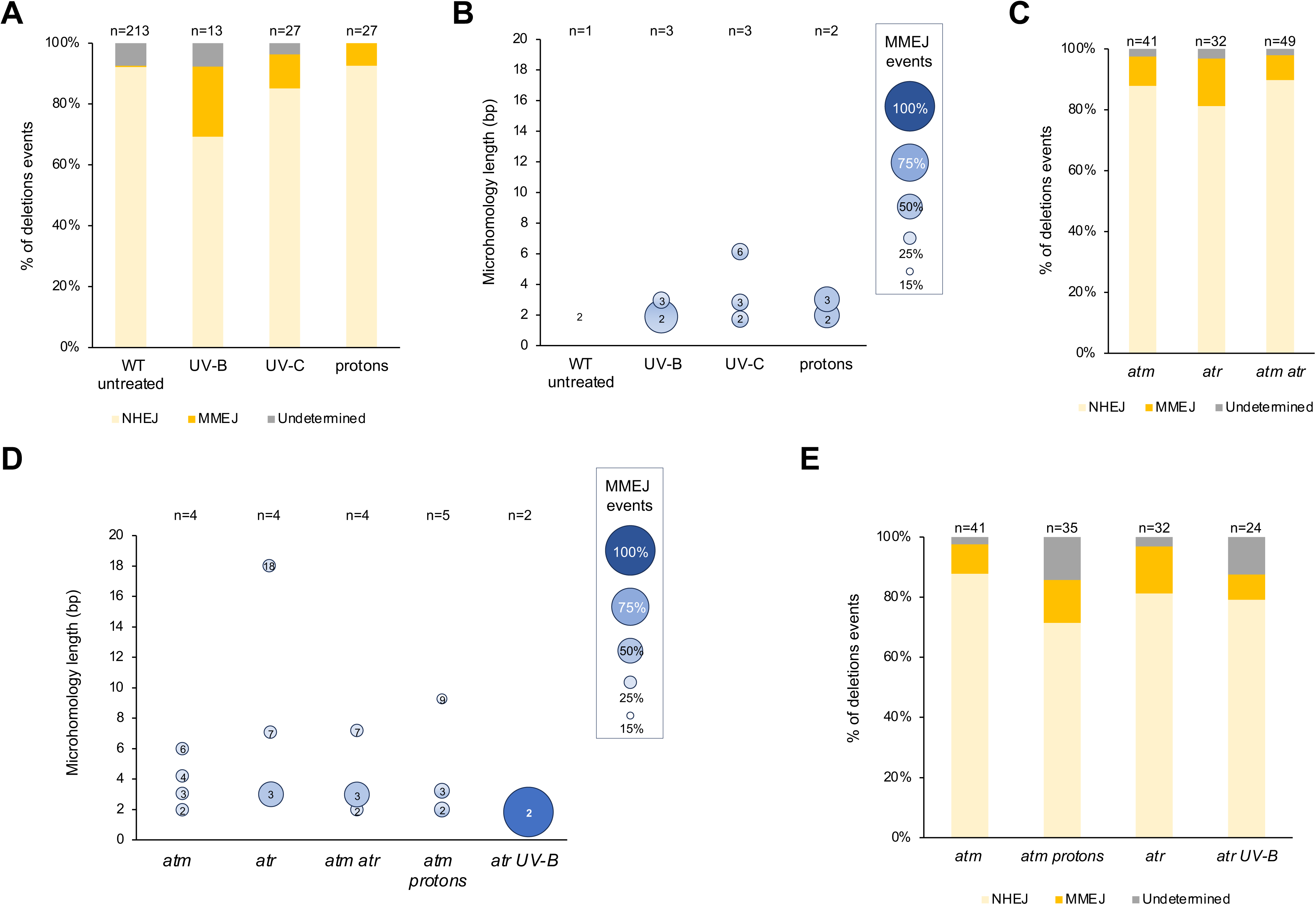
Characterization of end-joining repair mechanisms. **A.** Histogram representing the distribution of the NHEJ or MMEJ sequences signatures identified in the flanking regions of deletion in UV-B-, UV-C- and protons-treated WT Arabidopsis plants. n= total number of deletions. NHEJ: Non-Homologous End-Joining; MMEJ: Microhomology-Mediated End Joining; Undetermined: low sequence quality in one of the flanking regions. **B.** Bubble chart representing the microhomology length and their frequency within MMEJ events identified upon UV-B, UV-C and protons irradiation of WT Arabidopsis plants. n= total number of MMEJ events. Size of the microhomology is indicated in each bubble. **C.** Same as **A.** for *atm*, *atr* and *atm atr* plants. **D.** Same as **B.** for *atr*, UV-B irradiated *atr*, *atm* and protons-irradiated *atm* plants. **E.** Same as **A.** for *atm*, *atr*, *atm atr*, UV-B irradiated *atr* and protons-irradiated *atm* plants.

Single *atm* and *atr* mutant plants as well as double *atm atr* mutant plants exhibit between 8 and 15% of the sequences flanking directly the deletion with homologous repeats ranging from 2 to 18 bp in length, showing that MMEJ repair has been used (Fig. 8C, 8D and Table S1). A median effect could be observed in *atm atr* plants compared to each single mutant plants (Fig. 8C) suggesting that both PI3-like kinases might prevent independently the use of the MMEJ repair pathway.

Upon exposure to protons and to UV-B, *atm* and *atr* plants did not exhibit significant changes in the use of NHEJ and of MMEJ repair pathways (Fig. 8E). The length of the micro homologies remains unchanged in *atm* treated plants compared to untreated plants (Fig. 8D) suggesting that ATM plays a minor role in homology-directed repair in response to ionizing radiations. Conversely, in UV-B-irradiated *atr* plants, we identified a decrease of the number of MMEJ events and of the length of the micro-homologies (Fig. 8D). This is likely that UV-B exposure triggers the use of other repair pathways, that quantitatively and qualitatively affect the MMEJ mechanism.

These results provide an overview of the balance between repair mechanisms (NHEJ and homology-directed DSB repair), in WT and in DDR mutant plants, that have led to deletions upon different treatments.

## DISCUSSION

The use of long reads sequencing technology allowed characterizing the rearrangements of the linear genome structure of WT Arabidopsis plants and of DDR deficient plants (*atm* and *atr*) exposed to non-ionizing (UV-B and UV-C) and ionizing radiations (protons). We identified that most of the radiations-induced SVs occurred in heterochromatic regions in both WT and DDR deficient Arabidopsis plants. We found that the 2 PI3-like protein kinases, ATM and ATR, restrict INDELs formation. We determined that deletions are the outcome of NHEJ and homology-directed DSB repair pathways and identified that ATM and ATR repress the MMEJ repair pathway.

A crucial step in such comparative genomic approach was to quantify the putative variability that may contain plants originating from our collection of WT Col-0 Arabidopsis seeds in order to define our reference genome and to identify radiations-induced SVs. Interestingly, we found, in 3 independent biological replicates, more than 800 SVs compared to the TAIR10 reference genome with a core of 138 SVs. This common set of SVs reflects that long read sequencing technology allowed identifying slight differences between the TAIR10 reference genome and plants originating from our collection of WT (Col-0) Arabidopsis seeds. The higher resolution of the linear genome organization displayed by the long reads sequencing technology could explain the identification of these SV. Such improvement of the sequencing resolution was already validated with the Col-*CEN Arabidopsis thaliana* genome assembly that recently resolved all five centromeres (Naish et al., 2021). This holds true for the accurate assembly of others types repetitive sequences, TEs and small INDELS (Naish et al., 2021; Lian et al., 2024). Importantly, the SVs identified in WT Col-0 plants, may also have occurred during plant development in cells giving rise to the germline and/or during meiosis. In addition, the heterogeneity of the somatic plant cells (*i.e.*, leaves) could have contributed to the large number of SV found in the 3 independent biological replicates. Indeed, it has been reported that independent Arabidopsis reporter lines harboring a homologous recombination substrate exhibit different recombination rates in control conditions and also upon exposure to genotoxic stresses, highlighting that DSB formation and repair efficiency vary between genomic regions, plant material and plant species (Swodoba et al., 1994, Molinier et al., 2004). This could also be true for other types of repair processes and would lead to SV.

We identified that most of the radiations-induced SV occurred within constitutive heterochromatin suggesting that this part of the genome tends to be more prone to be rearranged than euchromatic regions (Lian et al., 2024). Mutation frequency was reported to be reduced in gene bodies and in essential genes (Monroe et al., 2022), highlighting that genic regions are more efficiently protected from different types of alterations. Thus, it strongly suggests that genomic and epigenomic features likely contribute to prevent deleterious mutations, rearrangement by either reducing DNA damageability and/or activating particular DNA repair pathways.

It was demonstrated that CRISPR–Cas9-induced SV as well as the repair outcomes are highly influenced by chromatin features (Filler-Hayut et al., 2021; Weiss et al., 2022). Repressive chromatin landscape, (*i.e.*, high DNA methylation), reduces CRISPR–Cas9 mutagenesis efficiency (Weiss et al., 2022). The discrepancy between the frequency of radiations- and CRISPR-Cas9-induced SVs in heterochromatin might reflect different damaging and/or repair efficacies in this part of the genome. Other features of constitutive heterochromatin could explain the occurrence of elevated frequency of radiations-induced SV. Constitutive heterochromatin contains high amounts of repeats that are source of homologies for repair (Avramova, 2002; Orel et al., 2003; Schmidt and Anderson, 2006). Indeed, DSB formed in repeated DNA sequences often leads to chromosomal rearrangements (Pâques et al., 1998). For example, centromeric regions containing 180 bp repeats, display high structural dynamics (Naish and Henderson, 2024). CNV has been characterized upon CRISPR-Cas9-induced DSBs at 45S rDNA and in plant deficient for WSS1A an important factor involved in the repair of DNA-protein crosslinks (Lopez et al., 2021; Hacker et al., 2022). Moreover, SVs in repeats are likely less deleterious than rearrangements occurring in single or low copies PCG. Our observation is in agreement with the recent study showing that the CRISPR-Cas9 induction of DSB in different tomato PCG leads predominantly to a precise repair (Ben-Tov et al., 2024). Thus, genome surveillance efficiency differs between euchromatin and heterochromatin, due to the presence of repetitive sequences and likely to others uncharacterized features.

The formation of higher amounts of DNA damage, the activation of specific DNA repair (sub)pathways or the presence of particular repair intermediates could also influence the formation of SV. Indeed, UV-induced photodamage are enriched in constitutive heterochromatin (Graindorge et al., 2019; Johann to Berens et al., 2023). These photolesions are predominantly processed by excision repair (*i.e,* Global Genome Repair; Molinier, 2017) which is a slow DNA repair mechanism that could favor the presence of repair intermediates that likely become substrates for rearrangements (Shärer, 2013). In euchromatin, hypo-mutation was demonstrated to be associated with H3K4me1-rich gene bodies and essential genes (Quiroz et al., 2024) whereas T-DNA and TE insertions occur preferentially in PCG and in their downstream regions, respectively (Brunaud et al., 2002; Sigman and Slotkin, 2016; Zhang et al., 2023,). Thus, euchromatin exhibits different behavior against point mutation and insertions, suggesting that the epigenome-recruitment of DNA repair factors is complex and tightly regulated regarding the type of induced DNA damage.

An emerging notion fed by different genome wide approaches highlighted that particular genomic regions are more prone to be rearranged. Indeed, hot spots of rearrangements (HOT; Jiao and Schneeberger, 2020) have been identified in Arabidopsis. We identified that more than 67% of the radiations-induced SV, overlap with HOT regions. These genomic regions have been proposed to undergo different evolutionary dynamics as compared to the rest of the genomes (Jiao and Schneeberger, 2020). The high occurrences of radiations-induced SVs in HOT regions, suggest that these regions still display flexibility and likely contain genomic and epigenomic features that favor rearrangements.

ATM and ATR are master regulators of the DDR (Shiloh, 2001). Long reads sequencing technology offered the possibility to study, with an improved resolution, the roles of ATM and ATR in the maintenance of genome integrity. We found that in absence of genotoxic stress each kinase restricts the formation of INDELs and that the number of SV did not significantly change in *atm atr* double mutant plants. These SV occurred predominantly in TE and IR, highlighting that despite the absence of these key kinases, PCG remained efficiently protected from rearrangements. Upon DSB formation, ATM phosphorylates the histone variants H2A.W.7 found in heterochromatin and H2A.X found in euchromatin (Lorković et al., 2017). Given that most of the SV identified in DDR deficient plants are located in heterochromatin it is likely that the absence or reduction of H2A.W.7 phosphorylation failed to efficiently mark DSB, inducing illegitimate recombination and chromosomal rearrangements. Such repair outcome would have also been expected in euchromatin with H2A.X, but genome integrity in PCG was properly maintained. This suggests that either DSB occur preferentially in heterochromatin or that other key regulatory factors trigger the DDR in genic regions in an ATM-/ATR-independent manner.

The analysis of both inserted and deleted regions allowed characterizing the origins of these sequences and the type of the repair pathway used, respectively. Transposition events (Mirouze et al., 2009; Debladis et al., 2017), extra chromosomal circular DNA (ecc DNA; Ito et al., 2011) and capture of DNA sequences from distant genomic regions (Gorbunova and Levy, 1997) can lead to neo insertions into the genome. For example, in *Arabidopsis thaliana* the *Ty1*/*Copia*-like retrotransposon, *ONSEN*, could be mobilized upon heat stress exposure (Ito et al., 2011) and *de novo* integrations of TE-derived eccDNA have been characterized in genic regions (Zhang et al., 2023). Such neo-insertions could lead to change in expression of neighboring PCG (Roquis et al., 2021; Thieme et al., 2022) and has been shown to be a major source of genetic variation in *A. thaliana* (Baduel et al., 2021). Thus, in plants exposed to non-ionizing and ionizing radiations, we expected to identify similar insertion events. However, we only found truncated TE, in both WT and DDR deficient plants.

During NHEJ repair genomic sequences of different lengths and distant from the break point can be copied by synthesis-dependent strand annealing like mechanisms (Rubin and Levy, 1997; Salomon and Puchta, 1998; Gorbunova and Levy, 1997; Gorbunova and Levy, 1999). Moreover, we found a preference for the intrachromosomal origin of the inserted sequences supporting the idea that genomic regions in the vicinity of the DNA breaks are used as template. This is likely that linear 1D organization and/or 3D genome folding favor such preferential use of intramolecular templates. Thus, these genomic features should be considered as parameter that could influence template availability during DSB repair.

Repeats/TE-(Cohen et al., 2008; Lanciano et al., 2017;) or intramolecular recombination-derived (Peterhans et al., 1990; Molinier et al., 2004) eccDNA are unstable episomes that can reintegrate and lead to SV (Peng et al., 2022). Arabidopsis plants with altered methylome and/or silencing machinery exhibited accumulation of TE-derived eccDNA and neo-insertions leading to SV (Zhang et al., 2023). Intramolecular recombination events produce ecc repair intermediates that have been shown to reintegrate into the genome and likely to form SV (Molinier et al., 2004). Thus, different origins of episomes exist and are source of genetic variability.

The loss of genetic information also occurs upon DSB repair. Indeed, deletions are the outcome of the NHEJ or of homology-directed repair pathways (Puchta 2005; Schubert and Vu, 2016). Nevertheless, it is important to notice that some NHEJ events can also be associated with filler DNA insertions (Gorbunova and Levy, 1997). Vu *et al*. (2017) showed that deletions occurred mainly via SSA and NHEJ. Importantly, repair events leading to deletions are mainly due to NHEJ if long homologous sequences are missing in the vicinity of the break. Nevertheless, short homologies (between 2 to 25 bp), could be used by the MMEJ pathway. The genomic sequences surrounding deletions in plants exposed to genotoxic stresses showed that NHEJ and MMEJ pathways have been preferentially used. NEHJ patterns represent the main outcome of repair and MMEJ events have been identified only in irradiated and in DDR deficient plants. Although, homology-directed repair pathways (SSA, MMEJ) have been described to be the main DSB repair pathways used in higher eukaryotes (Puchta, 2005) we did not identify canonical SSA events with 20 to 25 bp micro-homologies. This could be likely due the type of induced DNA damage and/or their genomic location. Indeed, the availability of particular repair factors could favor the use of one or others pathways as well as the nucleotide context (Ceccaldi et al., 2016). MMEJ acts in DSB repair in yeast, mammals (McVey and Lee, 2008) and plants (Heacock et al., 2004). Moreover, MMEJ seems to play an important role during the CRISPR/Cas-based genome editing (Ata et al., 2018; Tan et al., 2020; Weiss et al., 2020). A synthesis-dependent microhomology-mediated end joining (SD-MMEJ) mechanisms was also described and relies on *de novo* DNA synthesis to create microhomology (Yu and McVey, 2010). Plant organellar DNA polymerases have been shown to repair DSB by MMEJ (Garcia-Medel et al., 2019) suggesting that a SD-MMEJ-like mechanism exist in plants.

Using third generation sequencing technology we documented the effect of genotoxic stresses exposure on the whole linear genome structure shedding the light on the stability of euchromatin *vs* the pronounced flexibility of heterochromatin.

The experimental approach and the produced resources open new perspectives to further study the damaging effect of particular genotoxic agents and the type of DNA repair processes used within genome and epigenome complexity.

## EXPERIMENTAL PROCEDURES

### Plant material and growth conditions

*Arabidopsis thaliana* ecotype Col-0, was obtained from the Arabidopsis Biological Resource Stock Center (ABRC, Nottingham, UK). Plants were cultivated in soil in a culture chamber under a 16 h light (light intensity ∼150 μmol m^−2^ s^−1^; 21°C) and 8 h dark (19°C) photoperiod. *Arabidopsis thaliana atm-2* and *atr*-2 (Vespa et al., 2005) plants (Col-0 ecotype) were also used. The progeny of *ATM* +/-plants was genotyped to recover all *ATM* -/- plants used in this study. *ATR* -/- plants is the 3^rd^ generation of selfed *ATR* -/- plants. Double *atm atr* mutant plants were obtained by the crossing of *ATM* +/- plants with *ATR* -/- plants. F2 plants were genotyped to retrieve double *atm atr* mutant plants. Eight *atm atr* plants have been used for long reads sequencing experiments.

### Protons irradiation

Soil-gown 21-day-old Arabidopsis Col-0 plants (WT or *atm*) were exposed to 100 Gy of protons delivered by the TR24 cyclotron (https://cyrce.fr/en/home/) with a proton beam energy of 25 MeV (dose rate: 0.55 Gy/s). Leaves number 5 of 20 plants were irradiated on a zone of 5 mm of diameter. 24h upon exposure the irradiated zone was cut with a hollow punch (5 mm) and snap frozen in liquid nitrogen. Leaves discs of twice 20 untreated plants have been harvested from the 2 independent biological replicates and pooled. This untreated plant material corresponds to replicate 1 for the determination of our WT Col-0 pedigree.

### UV-B irradiation

Soil-gown 21-day-old Arabidopsis Col-0 plants (WT or *atr*) were exposed during 15 min to 4 bulbs of UV-B Broadband (Philips - TL 40W/12 RS SLV/25) to deliver a total dose of 6750 J/m^2^. Leaves number 5 of 10 irradiated plants have been harvested 24h upon irradiation and snap frozen in liquid nitrogen. Two independent biological replicates have been performed and pooled for long reads sequencing experiments. Leaves number 5 of twice 10 untreated plants have been harvested from the 2 independent biological and pooled. This untreated plant material corresponds to replicate 2 for the determination of our WT Col-0 pedigree.

### UV-C irradiation

Soil-gown 21-day-old Arabidopsis Col-0 plants were exposed to 2000 J/m^2^ of UV-C using the Stratalinker 2400 (Stratagene). Leaves 5 of 10 irradiated plants have been harvested 24h upon irradiation and snap frozen in liquid nitrogen. Two independent biological replicates have been performed and pooled for long reads sequencing experiments. Leaves number 5 of twice 10 untreated plants have been harvested from the 2 independent biological and pooled. This untreated plant material corresponds to replicate 3 for the determination of our WT Col-0 pedigree.

### Genomic DNA extraction and library preparation

Genomic DNA was prepared from 100 to 200 mg of leaves using the plant DNA extraction kit Nucleon Phytopure (Cytiva). RNaseA/T1treatment was performed and DNA was cleaned up using the MaXtract High Density kit (Qiagen) to recover High Molecular Weight DNA (HMW). Ultra-long DNA library for Nanopore sequencing was produced from 100 to 200 fmoles of HMW genomic DNA using the NEBNext companion module (NEB) and the Ligation Native Sequencing Kit V9 (ONT). 5– 50 fmoles of the library was loaded onto ONT FLO-MIN R9.4.1 or ONT FLO-PRO R9.4.1 R9.4.1 flow cells.

### Identification of genomic structural variants

Reads were sequenced on ONT FLO-MIN R9.4.1 or ONT FLO-PRO R9.4.1 R9.4.1 flow cells and basecalled with ont-guppy-gpu_6.3.8 with the model dna_r9.4.1_450bps_sup.cfg (Table S2). The analysis was performed using a Snakemake script adapted from the Oxford Nanopore Structural Variant pipeline (https://github.com/nanoporetech/pipeline-structural-variation). Sequencing quality was evaluated with MinIONQC (V 1.33.5). The mapping was performed with Minimap2 (V 2.17) using the *Arabidopsis thaliana*, Col0-TAIR10, as reference genome. The mapping quality was checked with Nanoplot (V 1.30.0). Coverage was evaluated by mosdepth (V 0.2.7) and Sniffles (V 1.0.11) was used to identify the structural variations (SV) in comparison with the reference genome Col0-TAIR10. SVs have been filtered using the following parameters: minimal SV length 1 pb, maximal SV length 1,000,000 pb, minimal read length 1,000 bp and minimal mapping quality 20. SV of the same type (insertion, deletion, duplication, inversion or inversion duplication) with the same genomic coordinates (Chr, start-end) ± 50 bp have been considered as identical.

### Characterization of the origins of insertions

Inserted sequences have been blasted using ncbi-blast+ (blastn fonction) and annotated to retrieve the origin of each insertion.

### Characterization of NHEJ and MMEJ repair events

The flanking sequences (± 50 bp) of the deleted regions have been fetched using samtools (command samtools faidx). Microhomologies have been manually curated to determine the rate of MMEJ events.

## AUTHOR CONTRIBUTIONS

SOS performed plant work and the bioinformatic analyses; AA performed the long reads sequencing; SS performed the plant work and the UV irradiations; SG set up the bioinformatic analysis; MP and JS performed the protons irradiations; QR and MR obtained the fundings, set up and performed the protons irradiations; JM designed experiments, obtained the fundings and wrote the manuscript.

## DATA AVAILABILITY

All sequencing data generated in this study have been deposited at SRA under the $$$$$$$$ repository. The fast5 files will be available on demand.

## ACKNOWLEDGEMENTS

This research was funded by the CNRS Mission for cross-cutting and interdisciplinary initiatives (MITI) in the frame of the program: Adaptation of the living to its environment program (AAP 2020), by the French National Research Agency (ANR-20-CE20-0021) and by the EPIPLANT Groupement de Recherche (CNRS, France). Salimata Ousmane Sall received a CNRS PhD grant (Doctorants CNRS 2020 Actions transverses en appui aux défis sociétaux).

## SUPPORTING INFORMATION

### SUPPLEMENTAL TABLES

**Table S1: MMEJ events**

**Table S2: Sequencing statistics**

### SUPPLEMENTAL FIGURES

**Supplemental figure 1:**
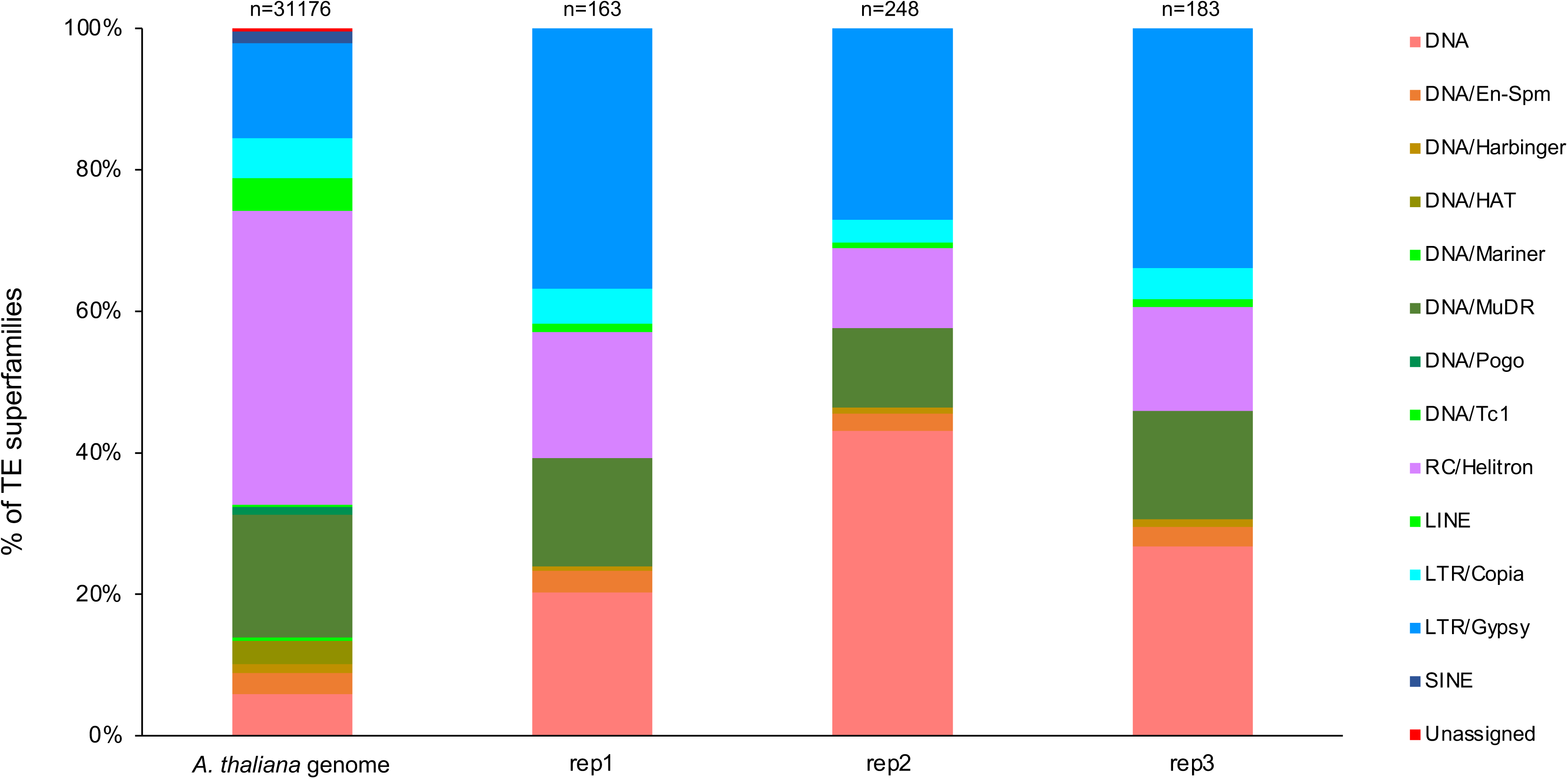
TE superfamilies exhibiting SV in WT arabidopsis plants. Histogram representing the distribution of TE superfamilies exhibiting SV identified in the 3 independent biological replicates of WT (Col-0) arabidopsis plants. Exact p values are shown (Chi square test).

**Supplemental figure 2:**
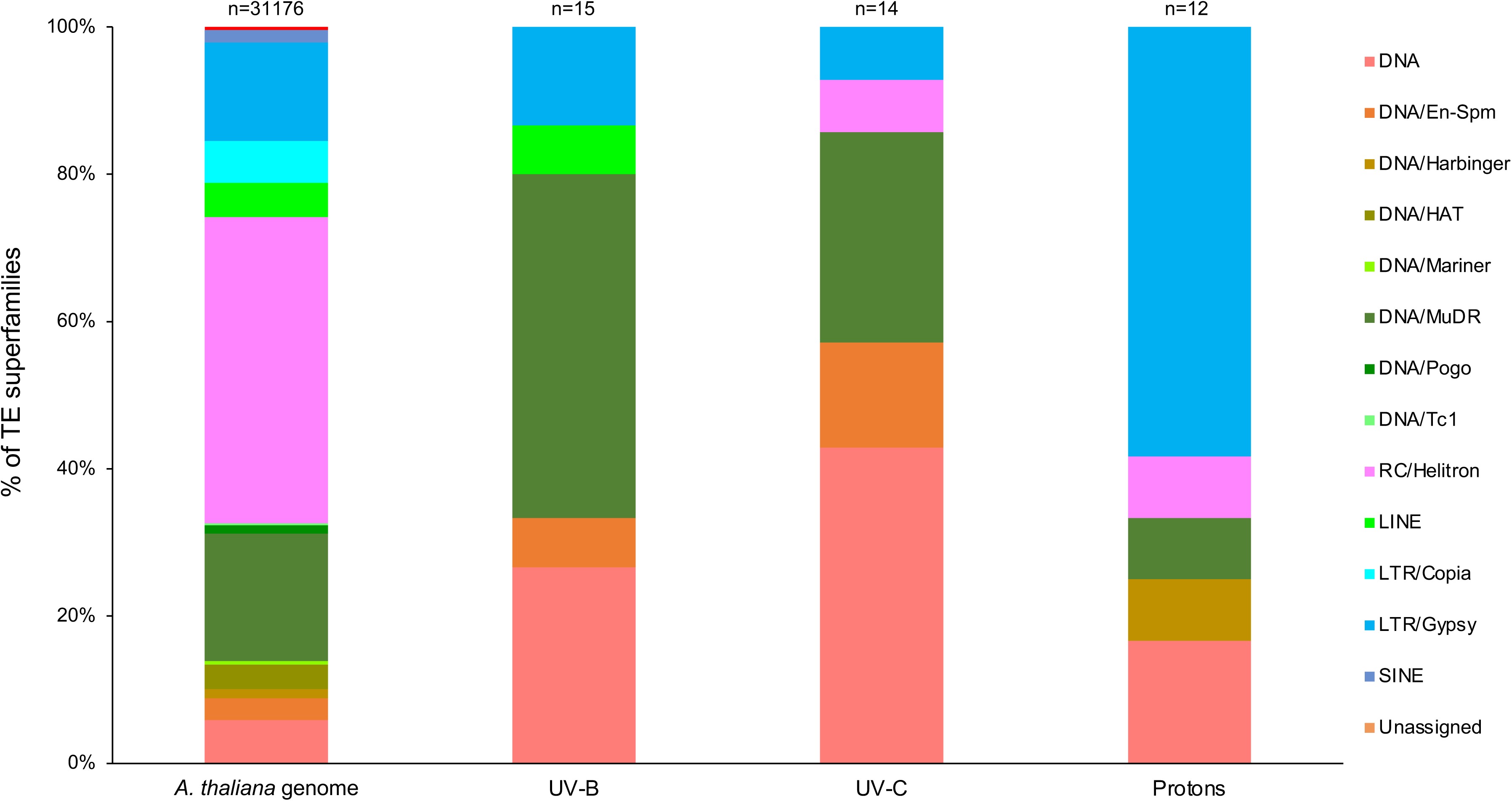
TE superfamilies exhibiting SV in radiation-treated WT arabidopsis plants. Histogram representing the distribution of TE superfamilies exhibiting SV identified in WT Arabidopsis plants treated with either UV-B, UV-C or protons. Exact p values are shown (Chi square test).

**Supplemental figure 3:**
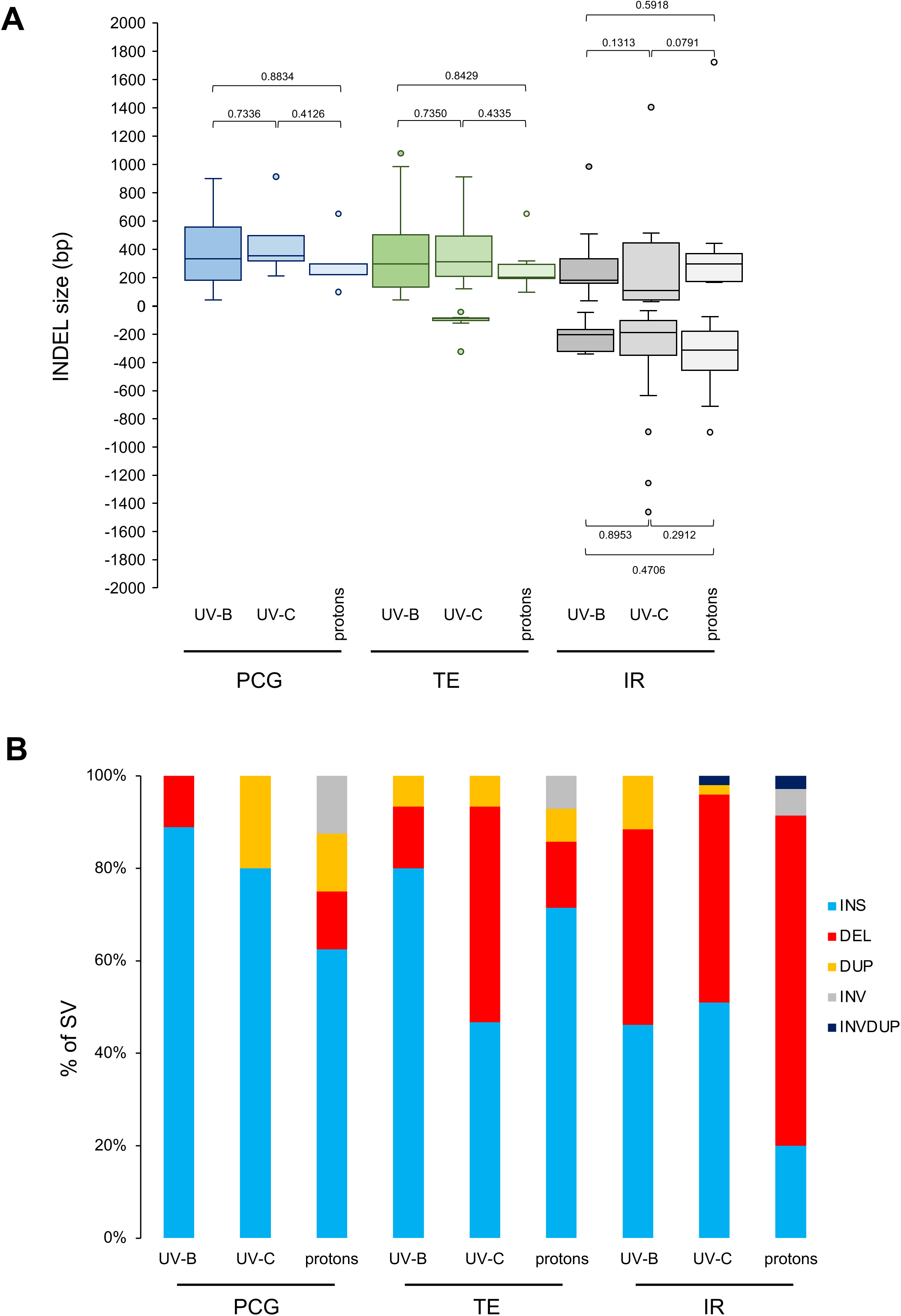
INDEL size in genetic elements of radiation-treated WT arabidopsis plants. **A.** Box plots representing the INDELs sizes identified in Protein Coding Genes: PCG; Transposable Elements: TE and Intergenic regions: IR. Exact p values are shown (Mann Whitney Wilcoxon test). **B.** Histogram representing the distribution of SV (INS: Insertion; DEL: Deletion; DUP: Duplication; INV: Inversion; INVDUP: Inversion Duplication) in PCG, TE and IR.

**Supplemental figure 4:**
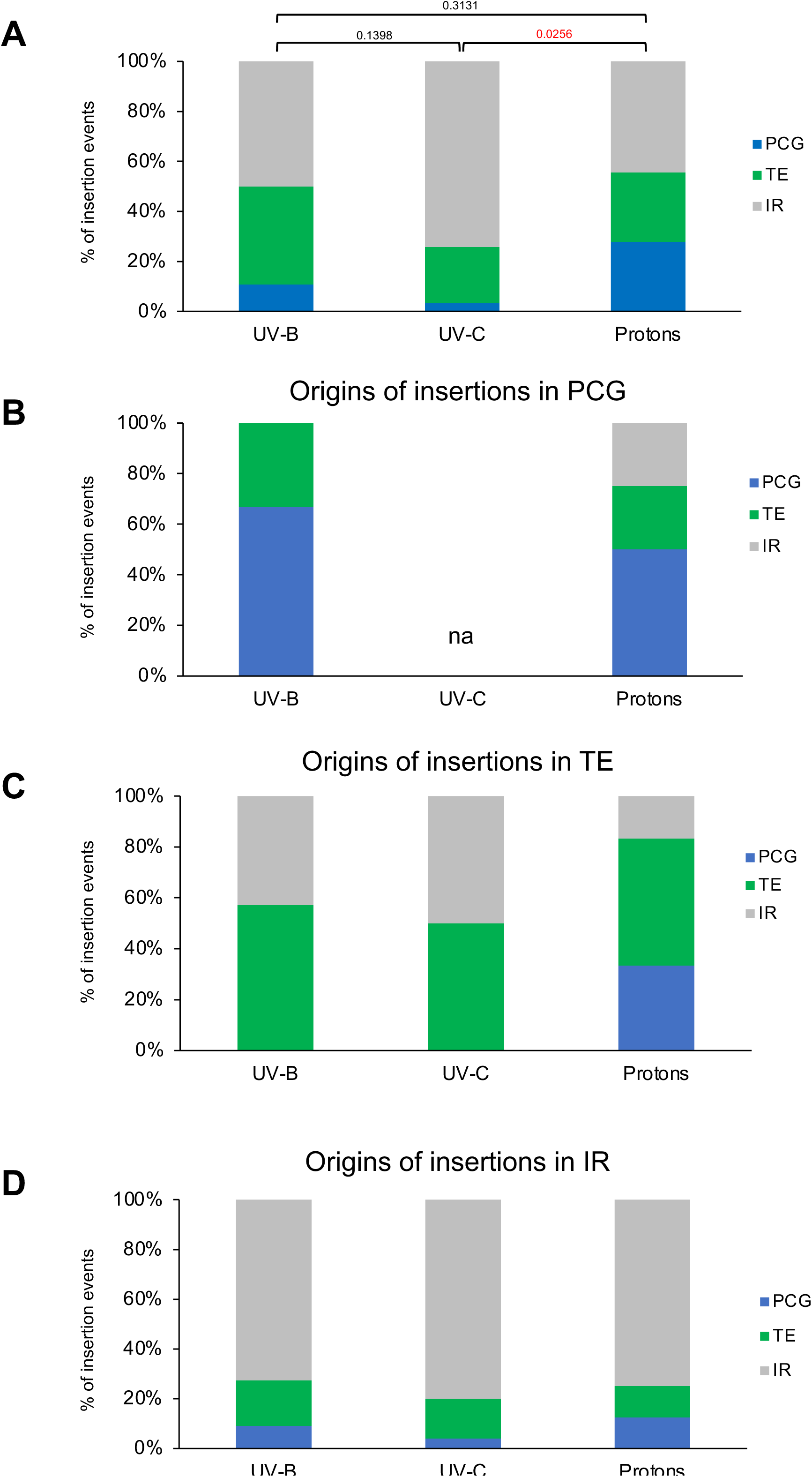
Origins of insertions in WT-irradiated Arabidopsis plants. **A.** Histogram representing the origin of the insertions (Protein Coding Genes: PCG; Transposable Elements: TE and Intergenic regions: IR) identified in WT Arabidopsis plants exposed to UV-B, UV-C or protons. **B.** Histogram representing the origin of the insertions identified in PCG of WT Arabidopsis plants exposed to UV-B, UV-C or protons. **C.** Same as **B.** for TE. **D.** Same as **B.** for IR.

**Supplemental figure 5:**
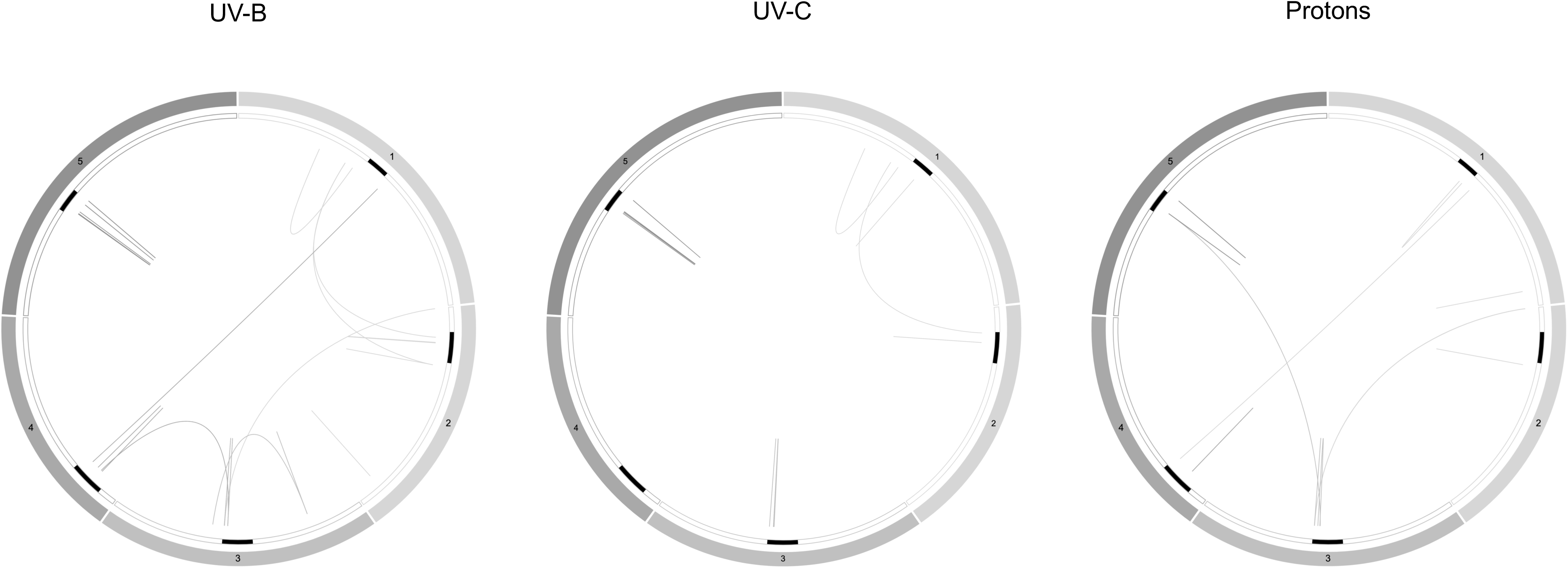
Origins of insertions in radiation-treated WT Arabidopsis plants. Circos representation of the origins of insertions in WT Arabidopsis plants treated with either UV-B, UV-C or protons. Black rectangles represent the centromeres.

**Supplemental figure 6:**
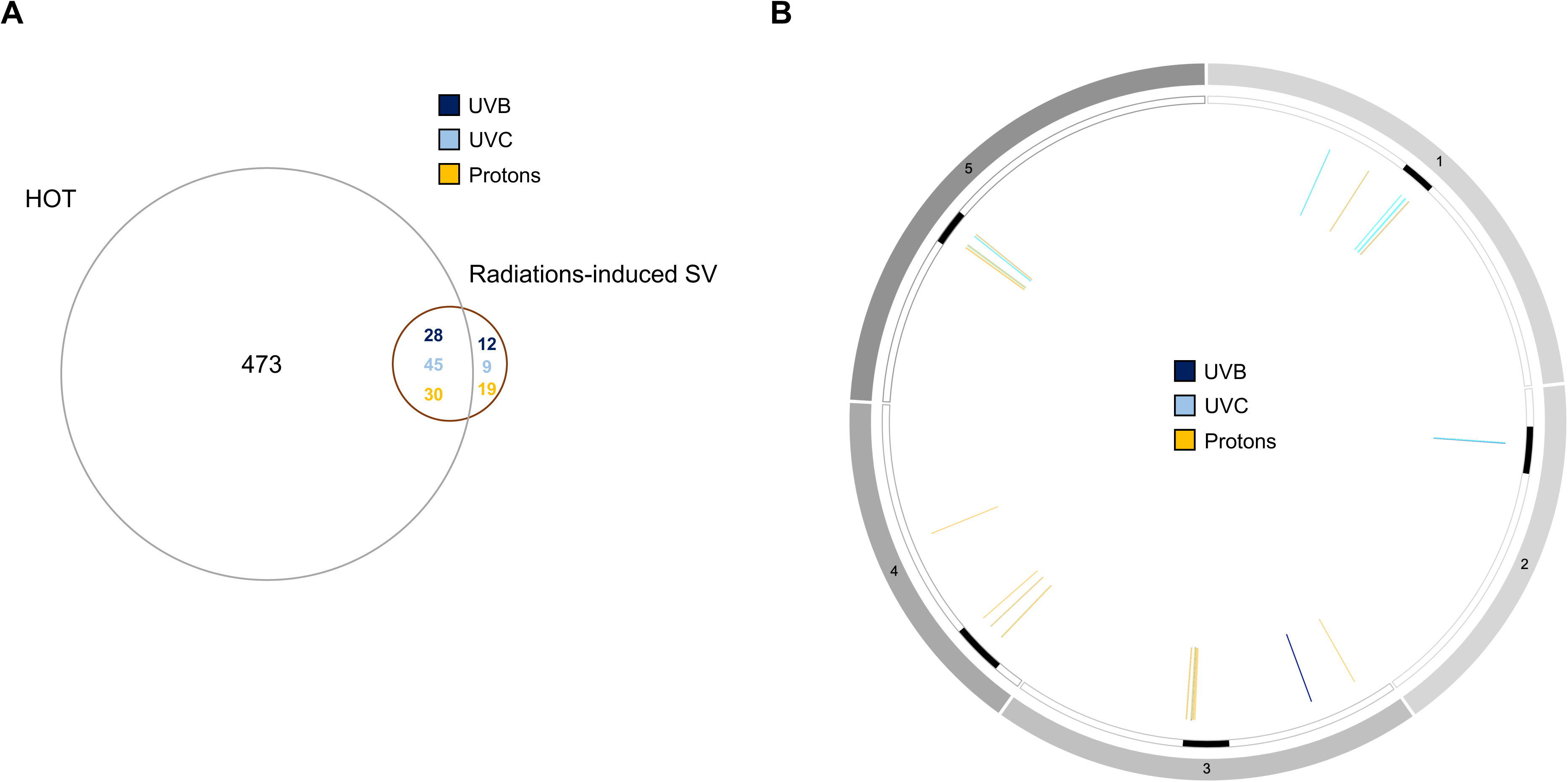
Genomic SV overlapping with HOT regions. **A.** Venn diagram representing the overlap of SV identified in irradiated WT Arabidopsis plants and hotspots of rearrangements (HOT; Jiao and Schneeberger, 2020). **B.** Circos representation of genomic SV of irradiated WT Arabidopsis plants overlapping with HOT. Black rectangles represent the centromeres.

**Supplemental figure 7:**
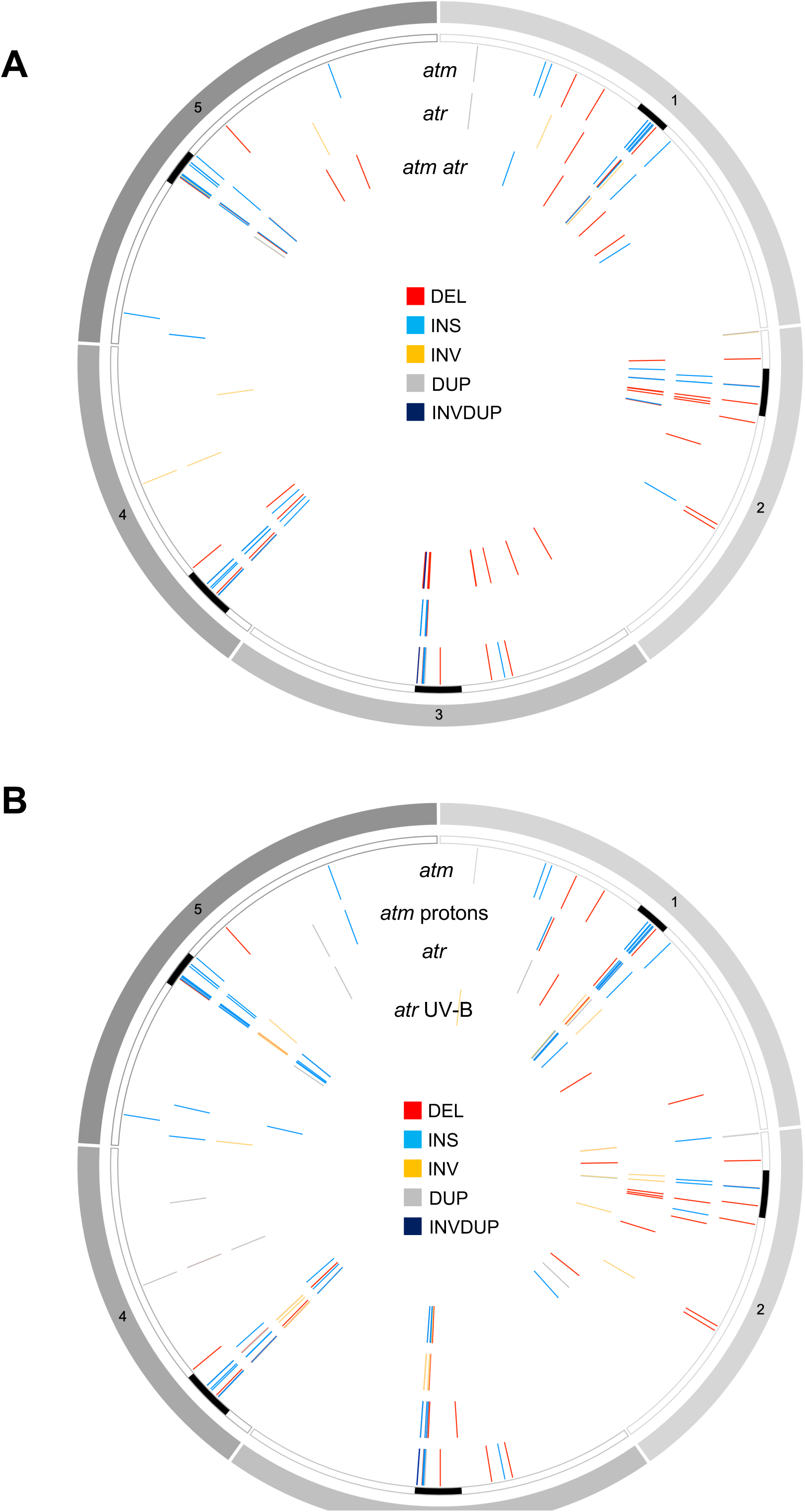
Genomic locations of structural variations. A. Circos representation of genomic SV (INS: Insertion; DEL: Deletion; DUP: Duplication; INV: Inversion; INVDUP: Inversion Duplication) identified in *atm*, *atr* and *atm atr* plants. **B.** Same as **A.** for *atr*, UV-B irradiated *atr*, *atm* and protons-irradiated *atm* plants. Black rectangles represent the centromeres.

**Supplemental figure 8:**
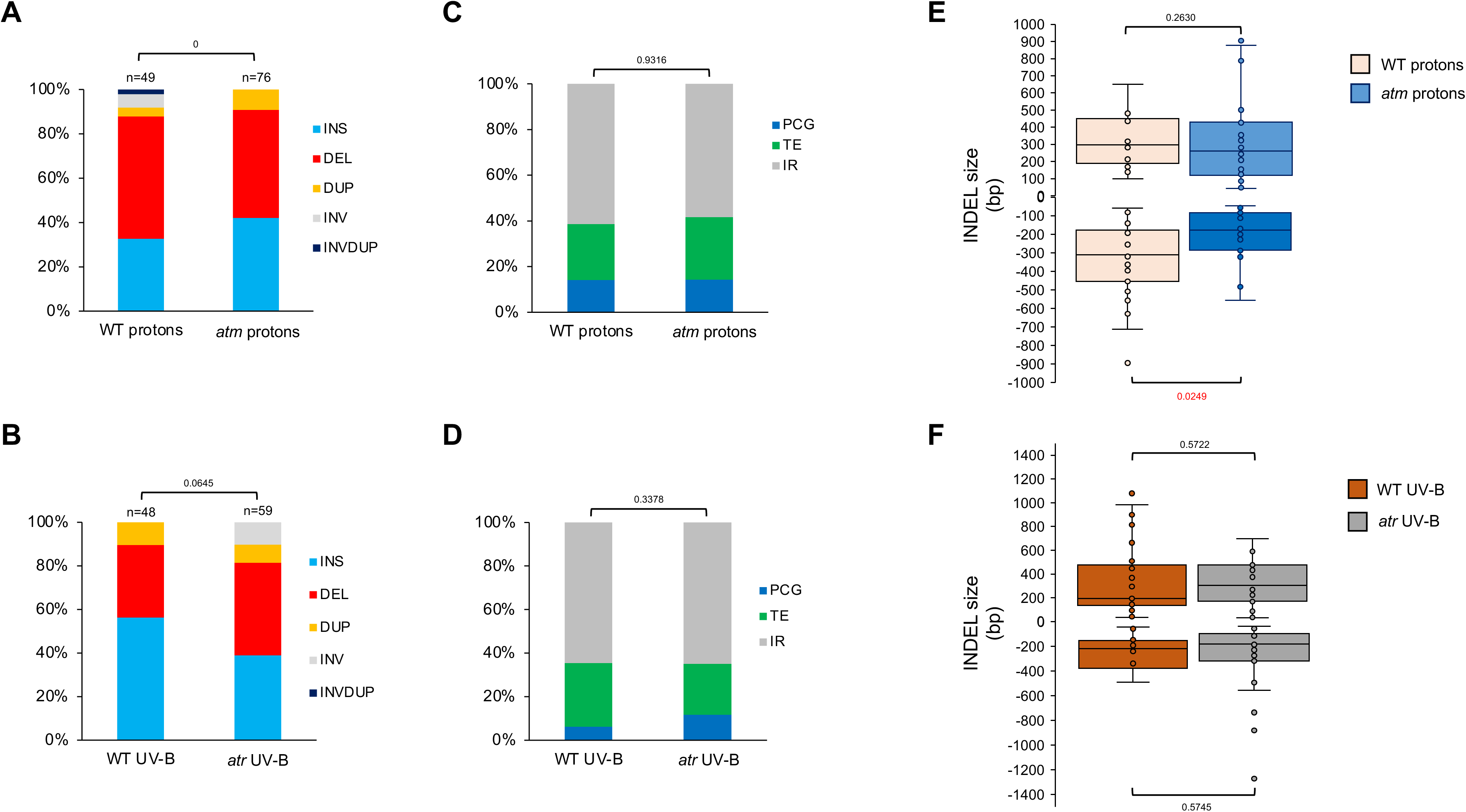
Comparisons of the radiation-induced genomic structural variations between WT, *atr* and *atm* Arabidopsis plants. **A.** Histogram representing the distribution of the different types of genomic SV identified in WT and *atm* plants irradiated with protons. INS: Insertion; DEL: Deletion; DUP: Duplication; INV: Inversion; INVDUP: Inversion Duplication. n= total number of SV. Exact p values are shown (Chi square test). **B.** Same as A. for WT and *atr* plants irradiated with UV-B. **C.** Histogram representing the distribution of the genetic elements (Protein Coding Genes: PCG; Transposable Elements: TE and Intergenic regions: IR) exhibiting SV in WT and *atm* plants irradiated with protons. Exact p values are shown (Chi square test). **D.** Same as C. for WT and *atr* plants irradiated with UV-B. **E.** Box plots representing the size of the INDELs identified in WT and *atm* plants irradiated with protons. Exact p values are shown (Mann Whitney Wilcoxon test). **F.** Same as E. for WT and *atr* plants irradiated with UV-B.

**Supplemental Table 1:**
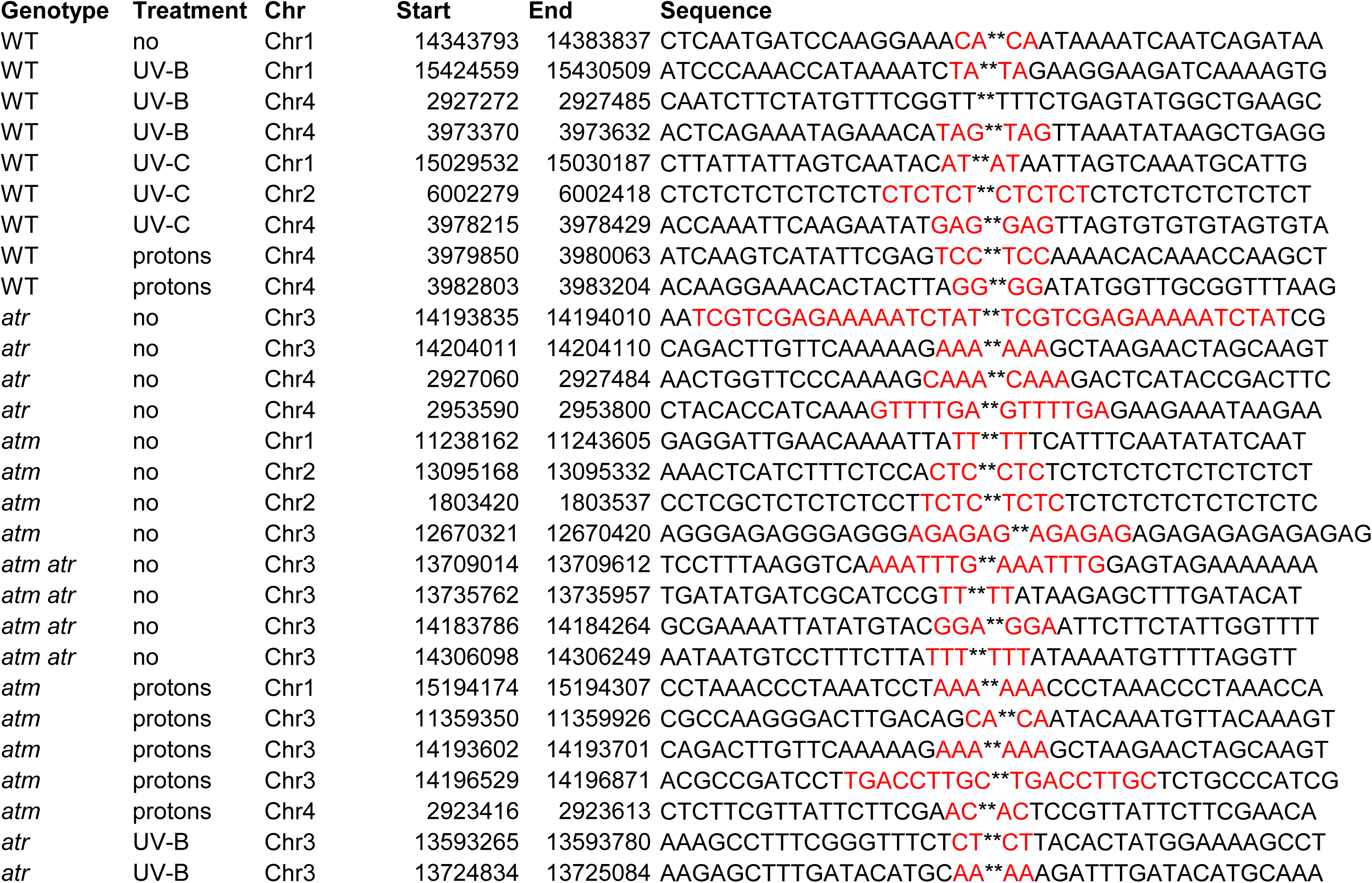
Nucleotides sequences of the genomic regions flanking deletions events and repaired by the MMEJ pathway. Microhomologies are shown in red

**Supplemental Table 2:** Sequencing statistics

| Sample | Genotype | Treatment | Mean<br>read<br>length<br>(bp) | Mean<br>read<br>quality | Median<br>read<br>Length<br>(bp) | Median<br>read<br>quality | Read<br>length<br>N50 kb | Total<br>Gb | Number<br>of reads<br>(x 1,000) | Mapped<br>fraction | Average<br>coverage |
| --- | --- | --- | --- | --- | --- | --- | --- | --- | --- | --- | --- |
| **WT rep1** | WT | no | 4,754.8 | 9.7 | 1,007 | 10.1 | 25.62 | 5.00 | 1,152.94 | 0.63 | 22.89 |
| **WT rep2** | WT | no | 4,073.9 | 13.6 | 884 | 14.2 | 19.93 | 10.53 | 2,900.00 | 0.82 | 60.56 |
| **WT rep3** | WT | no | 5,555.6 | 13.1 | 856 | 13.7 | 25.14 | 5.04 | 908.22 | 0.81 | 28.63 |
| **WT UV-B** | WT | UV-B | 3,160.7 | 9.6 | 1,028 | 10 | 11.18 | 10.38 | 3,290.00 | 0.88 | 55.62 |
| **WT UV-C** | WT | UV-C | 5,524.2 | 9.8 | 1,485 | 10.1 | 21.24 | 10.47 | 1,895.53 | 0.86 | 60.80 |
| **WT Protons** | WT | Protons | 5,462.3 | 9.9 | 1,151 | 10.3 | 21.90 | 3.87 | 4,098.09 | 0.90 | 51.88 |
| *atm* | *atm* | no | 6,432.5 | 9.7 | 1,052 | 10.1 | 26.87 | 2.10 | 2,485.28 | 0.50 | 12.90 |
| *atr* | *atr* | no | 3,724.7 | 9.1 | 714 | 9.5 | 23.23 | 13.92 | 3,883.81 | 0.41 | 76.75 |
| *atm atr* | *atm atr* | no | 8,754.7 | 12.6 | 5,965 | 12.4 | 14.00 | 11.88 | 861.426 | 0.93 | 41.45 |
| *atm* Protons | *atm* | Protons | 5,804.1 | 9.2 | 1,079 | 9.6 | 30.58 | 10.95 | 2,061.43 | 0.63 | 66.50 |
| *atr* UV-B | *atr* | UV-B | 5,702.7 | 9.1 | 905 | 9.5 | 29.48 | 10.97 | 2,139.53 | 0.58 | 66.38 |

